# Orthrus: Towards Evolutionary and Functional RNA Foundation Models

**DOI:** 10.1101/2024.10.10.617658

**Authors:** Philip Fradkin, Ruian Shi, Taykhoom Dalal, Keren Isaev, Brendan J. Frey, Leo J. Lee, Quaid Morris, Bo Wang

## Abstract

In the face of rapidly accumulating genomic data, our ability to accurately predict key mature RNA properties that underlie transcript function and regulation remains limited. Pre-trained genomic foundation models offer an avenue to adapt learned RNA representations to biological prediction tasks. However, existing genomic foundation models are trained using strategies borrowed from textual domains that do not leverage biological domain knowledge. Here, we introduce Orthrus, a Mamba-based mature RNA foundation model pre-trained using a novel self-supervised contrastive learning objective with biological augmentations. Orthrus is trained by maximizing embedding similarity between curated pairs of RNA transcripts, where pairs are formed from splice isoforms of 10 model organisms and transcripts from orthologous genes in 400+ mammalian species from the Zoonomia Project. This training objective results in a latent representation that clusters RNA sequences with functional and evolutionary similarities. We find that the generalized mature RNA isoform representations learned by Orthrus significantly outperform genomic foundation models on mRNA property prediction tasks, and requires only a fraction of fine-tuning data to do so. Finally, we show that Orthrus is capable of capturing divergent biological function of individual transcript isoforms.

## Main

Mature messenger RNAs (mRNA), resulting from transcription and splicing of precursor RNAs, encode essential genetic information for protein synthesis. The properties and functions of RNAs are often tightly linked to their sequence and are critical in modulating protein expression and cellular processes [1]. Experimental procedures [2–4] have been pivotal in studying RNA metabolism, but these techniques are often time-consuming and expensive. As an alternative, supervised machine learning models trained on genetic sequences can learn predictive features directly from RNA sequences, offering effective and low-cost prediction of post-transcriptional processes such as alternative splicing and RNA degradation [5–7]. These models can be used to identify disease mechanisms [8, 9], improve therapeutics such as mRNA vaccines [10], and predict the effects of perturbations [11]. Despite the importance of these applications, the difficulty associated with experimental acquisition of training data restricts the use of supervised methods for a wider range of tasks.

Recent works [12–19] proposed foundation models as an alternative to supervised learning approaches in genomic domains. Genomic foundation models use deep neural networks to learn an expressive representation of genetic sequences by pre-training on large datasets. During pre-training, self-supervised learning (SSL) objectives are used to train the model in the absence of labeled examples. SSL can be formulated through a data reconstruction objective, where a model is required to reconstruct a portion of the input data. Existing genomic foundation models use training objectives including next token prediction (NTP) and masked language modeling (MLM) [20, 21]. Foundation models that effectively capture the underlying biological complexities can enable property prediction and few-shot learning, generalizing to new experimental biology, such as mRNA half-life or RNA localization prediction, using a minimal number of samples [22]. However, current genomic foundation models, which primarily adapt self-supervised learning objectives from natural language processing, have struggled to fully realize this potential. The unique properties inherent to genomic data pose challenges for implementing reconstruction-based SSL objectives or supervised learning approaches.

Genomic sequences in the natural world are constrained by evolutionary viability, resulting in low natural diversity^1^ and high mutual information across genomes from the same species [24]. Approximately 10% of the human genome is under constraint and can be considered high information content [25, 26]. The remaining 90% of the genetic sequence lacks evidence of negative selection, meaning mutations may have little to no impact on organism fitness [27, 28]. Without a strong biological inductive bias, existing reconstruction-based SSL models often reconstruct non-informative tokens, which can result in sub-optimal representations. Due to the high-mutual information between samples, it is also difficult to scale the effective size of the training dataset to circumvent this issue. As such, direct application of these SSL methods to genomics has yielded representations that are often sub-optimal for mRNA property prediction tasks [12–14, 29]. Although impressive performance has been achieved by scaling some models to billions of parameters, these gains are not always proportional to the immense increase in computational cost. This trend of diminishing returns suggests that relying on scale alone may be an inefficient strategy for advancing mRNA representation learning.

Here, we propose Orthrus, an RNA foundation model that is pre-trained on mature RNA sequences. Orthrus uses a novel biologically motivated contrastive learning objective to structure the model latent space by maximizing similarity between splicing isoforms and evolutionary related transcripts [30, 31]. Using this contrastive objective, Orthrus is pre-trained on splicing annotation data from 10 species and orthologous alignments from more than 400 mammalian species in Project Zoonomia [32]. Pre-training Orthrus on mature RNAs with high functional importance and sequence conservation further allows Orthrus to focus on sequence regions with high information content [25, 27]. This allows Orthrus to be highly efficient, achieving state-of-the-art performance with orders of magnitude fewer parameters than many contemporary genomic foundation models. Orthrus is trained using a Mamba encoder, which enables favorable model properties such as the learning of variable motif spacing, context filtration, and linear memory scaling with sequence length [33]. Orthrus pre-training results in effective mature RNA representations that are predictive of diverse mRNA properties and functions.

We show that Orthrus’s learned representations can be used to accurately predict the properties and functions of mature mammalian RNA sequences in three key contexts. First, we test the effectiveness of biologically inspired contrastive learning by fitting a linear model on top of the pretrained latent representations. Orthrus outperforms most other self-supervised foundation models, and applying this simple linear transformation exceeds the performance of supervised methods on all property prediction tasks. Second, we fine-tune the pre-trained models on experimentally collected mRNA property datasets and demonstrate state-of-the-art performance when generalizing to unseen sequences. Orthrus is able to effectively perform in the low data regime, requiring as few as 30 labeled examples to fine-tune an mRNA half-life predictor. Finally, we demonstrate that Orthrus is capable of grouping RNA isoforms by their common functions in the model latent space. This brings us closer to being able to annotate the function of individual splice isoforms, a long-standing challenge in RNA biology.

## Results

### Biologically Inspired Contrastive Learning Dataset

Orthrus is trained using contrastive learning, which constructs a structured representation space by directly maximizing the embedding similarity within positive pairs of related RNA transcripts while minimizing the similarity of all unrelated transcripts. Each positive contrastive pair consists of a reference RNA transcript that is paired with a transcript sampled from a set of related mature RNA sequences. Under the contrastive learning framework, these related mature RNAs can be viewed as augmentations of the reference transcript, as they share similarities in function and property but differ in sequence. We construct a contrastive learning dataset that identifies positive pairs based on alternative splicing and orthologous transcripts produced through mammalian speciation events. Our approach is founded on the biological principle that mRNA sequences produced through these processes are more functionally similar to each other than to randomly sampled RNA transcripts [34–36].

To construct positive pairs based on alternative splicing, we group alternatively spliced transcripts using the GENCODE and RefSeq [38, 39]. We use splice information across 10 metazoan organisms: human, mouse, chicken, C. elegans, chimpanzee, cow, dog, fruit fly, rat, and zebrafish (Figure 1 A). While alternatively spliced mRNA isoforms exhibit sequence variability and can sometimes acquire novel functions, our approach is based on the principle that, isoforms on average are more functionally similar to each other than to randomly sampled transcripts [34, 35]. We empirically find that sequence diversity arising from different exon combinations in alternatively spliced isoforms serves as an effective source of function-preserving variation.

**Fig. 1:**
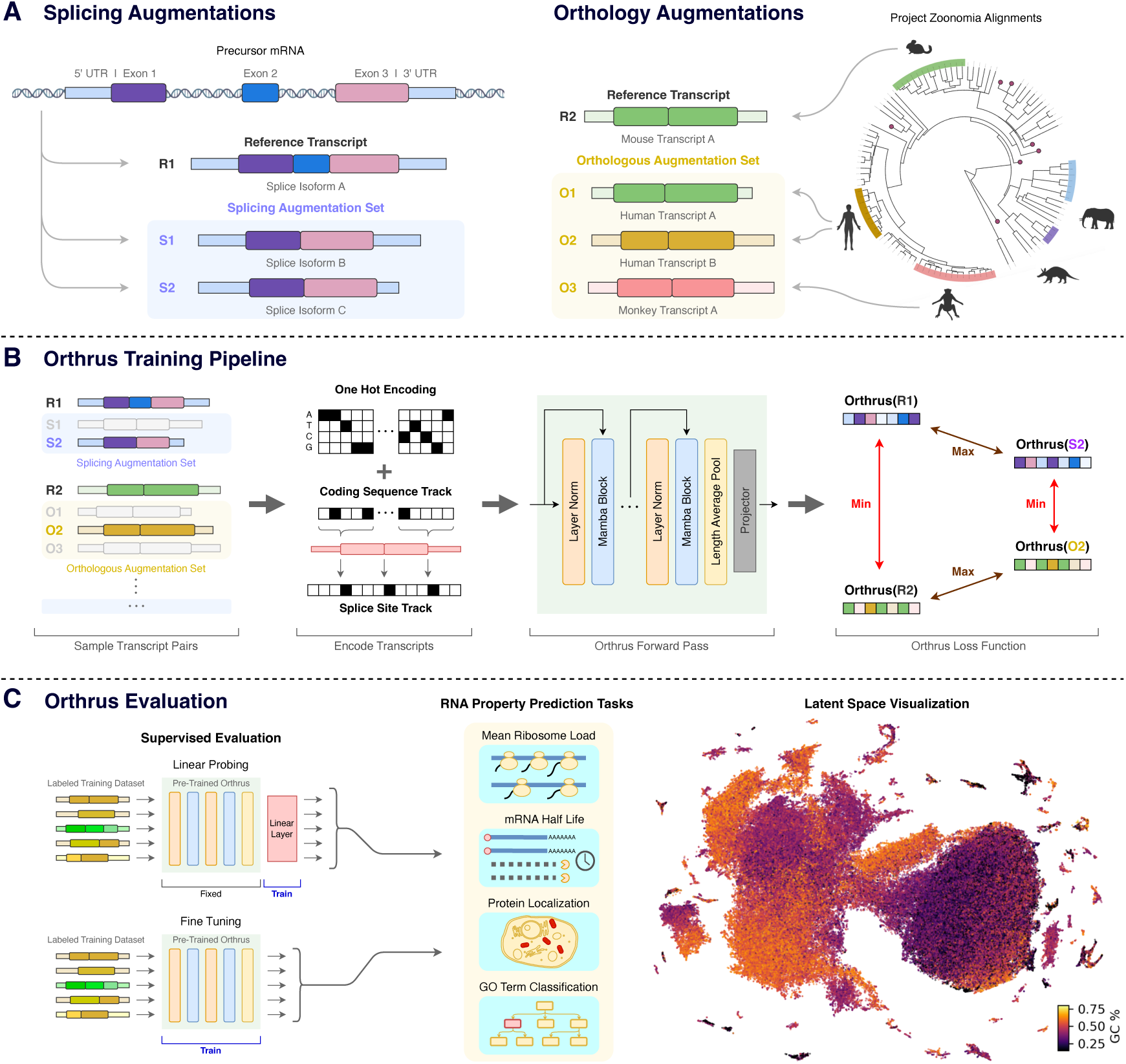
Overview of Orthrus. (A) Contrastive dataset construction: We treat each RNA transcript in the pre-training dataset as a reference transcript. For each reference transcript, we identify sets of transcripts that are related through alternative splicing and orthology. Each reference transcript can be associated to both splicing and orthology augmentation sets. **(B) Training pipeline:** For all reference transcripts in a batch, we randomly sample a positive paired transcript from its splicing and orthology augmentation sets. All transcripts are converted into a six track encoding. We then generate a projection of the sequences using Orthrus model and apply the contrastive loss over these samples, maximizing similarity between positive pairs while minimizing it for all the other transcripts. **(C) Evaluation:** Orthrus is evaluated in linear probing and fine-tuning contexts for a variety of mature mRNA property prediction tasks. Orthrus learns the structural properties of RNA transcripts, as shown by latent space visualization using UMAP [37].

Orthologous transcripts from mammalian species present another source of function-preserving sequence diversity [40, 41]. We use positive pairs generated through the process of speciation across the Eutheria clade derived from the Zoonomia TOGA resource [32], which performs joint gene annotation and orthology inference mapping transcripts from over 400 species to human and mouse annotations. To identify orthologous pairs, TOGA performs alignment over identified coding sequences and neighboring intronic and intergenic regions. Importantly, this exposes the model to transcript regions that are conserved over evolutionary time due to negative selection. These regions are likely to be functionally important and relevant for mRNA metabolism.

Overall, our final dataset contains 32 million unique transcripts and over 887 million unique positive pairs (Table 1, Section 7).

**Table 1:** Overview of contrastive datasets.

| Contrastive Datasets | Zoonomia | Splicing | # of Pairs | # of Transcripts |
| --- | --- | --- | --- | --- |
| Zoonomia Eutheria & Splicing Gencode Basic | ✓ | ✓ | 887,025,349 | 32,714,651 |
| Zoonomia Eutheria | ✓ | ✗ | 31,961,363 | 32,714,651 |
| Splicing Gencode Basic | ✗ | ✓ | 18,120,876 | 753,288 |
| No Augmentations | ✗ | ✗ | 0 | 753,288 |

**Table 2:** Overview of evaluation datasets.

| Task Dataset | Category | Number of Unique Sequences | Maximum Sequence Length | Homology Split Possible | Species |
| --- | --- | --- | --- | --- | --- |
| RNA Half Life Human | Regression | 12,968 | 12,288 | ✓ | Human |
| RNA Half Life Mouse | Regression | 13,738 | 12,288 | ✓ | Mouse |
| Mean Ribosome Load | Regression | 11,693 | 12,275 | ✓ | Human |
| Protein Localization | Classification | 9,769 | 12,275 | ✓ | Human |
| Gene Ontology MF | Classification | 5,737 | 12,236 | ✓ | Human |
| eCLIP | Classification | 21,559 | 205,012 | ✓ | Human |
| RNA-Localization Fazal | Classification | 3,297 | 14,897 | ✓ | Human |
| RNA-Localization Ietswaart | Classification | 10,043 | 13,459 | ✓ | Human |

### Orthrus Model Overview

During the contrastive training phase, we sample positive pair sequences from mature RNA transcript sets and maximize their similarity in the model latent space (Figure 1 B). Given a batch of *N* reference sequences, *x*^1^*, . . ., x*^1^, we construct positive pairs (*x*^1^, *x*^2^) by randomly sampling *x*^2^ from the augmentation set of transcripts related to *x*^1^ through alternative splicing or orthology processes, as described in the previous section. The augmentation set associated with each reference transcript can contain both splice isoforms and orthologous transcripts. The positive pair transcript from the augmentation set (*x*^2^) is re-sampled for each training epoch. We pass these positive pairs through a Mamba [33] encoder (*f_θ_*) resulting in the outputs *h*^1^ and *h*^2^. These representations are then fed into a multi-layer perceptron projection head (*g_θ_*) and the outputs are used to calculate normalized projections *z_i_*. We use the decoupled contrastive learning (DCL) loss [42] to perform the contrastive learning objective, pushing apart unpaired transcripts and maximizing the cosine similarity between positive pairs (Figure 1 B). After pre-training, we discard the projection head and directly use the outputs from the Mamba encoder as embeddings. The contrastive learning objective can be further combined with other self-supervised learning techniques such as masked language modeling objective (MLM). We introduce three versions of Orthrus using a backbone Mamba encoder: Orthrus Small consisting of 1.3 million trainable parameters, Orthrus with 10.1 million trainable parameters, and Orthrus MLM a 10.1 million parameter model which is trained with a combination of contrastive learning and masked language modeling objectives. Model hyperparameters are reported in Appendix A.3.

### Orthrus Embeddings are Predictive of Diverse mRNA Properties

To evaluate the effectiveness of our pre-trained representations, we followed the conventional evaluation strategy of linear probing. The learned latent embedding is effective if ∃ **w** s.t. **w***^T^* **X** + *b* = *y*^, where, **X** is a matrix of embeddings, and *y*^ are the predictions for labels *y*. To evaluate the above, we freeze the weights of the Mamba encoder *f_θ_* and train a linear layer to predict labels for regression and classification tasks. Further experimental details are described in Appendix A.4.

We quantitatively evaluate whether Orthrus embeddings contain information regarding key transcript properties such as UTR length, number of exons, CDS length, and transcript type in Figure 2 A. We observe that Orthrus embeddings are highly predictive of these attributes, which are important for predicting properties such as mRNA half-life [7]. We note that Orthrus embeddings are of a fixed length, meaning these properties cannot be simply derived as a function of embedding dimensionality. In Figure 2 B, we demonstrate that Orthrus excells at mRNA property prediction and outperforms other self-supervised methods on a diverse set prediction tasks. In particular, Orthrus outperforms or matches Evo2 [19] on seven tasks, despite having 700 × fewer parameters, highlighting the efficiency of its biologically grounded training objective. To establish a strong performance ceiling, we also train supervised *Ab initio* models for each task using the full set of ground-truth labels.

**Fig. 2:**
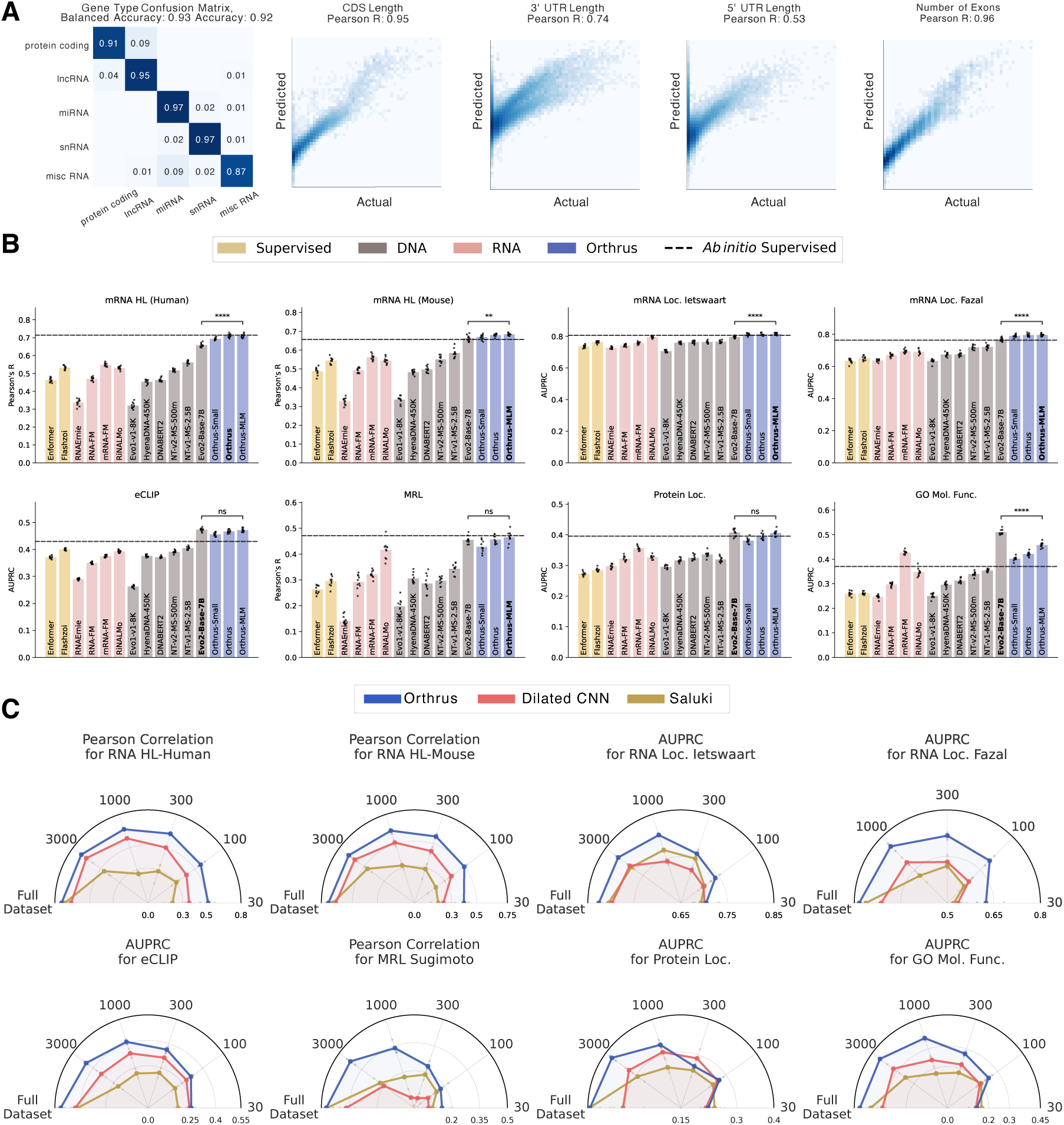
mRNA Property Prediction using Orthrus. **(A)** Evaluation of Orthrus’s latent representation by fitting a linear model to predict structural properties. The confusion matrix evaluates Orthrus’s ability to classify transcript types using logistic regression on learned embeddings. The four scatter plots assess Orthrus’s ability to predict structural RNA properties, including CDS length, 3’ UTR length, 5’ UTR length, and number of exons. **(B)** Benchmarking linear probing performance on mRNA property prediction tasks for self-supervised genomic foundation models. Error bars show 95% confidence intervals, constructed using 10 runs with homology aware data splits. The grey dashed line indicates performance of the maximum between fully fine-tuned Dilated CNN models and Saluki *Ab initio* models. Statistical comparison is done using a two sided t-test between the best Orthrus variant and the top performing non-Orthrus model. **(C)** Plots evaluating the fine-tuning performance of Orthrus Small across varying levels of data availability. Each dataset is subsampled to the indicated number of points on the circumference of the plot. Point estimates are plotted, averaged across three random seeds and random data splits.

For each task, we establish this baseline by selecting the stronger of two supervised architectures: the CNN-RNN hybrid from Saluki [7] or a dilated convolutional neural network [43], and indicate its performance with a dashed line in Figure 2 B. For all evaluated tasks, a linear model trained on Orthrus embeddings outperforms or matches the corresponding specialized, *Ab initio* baseline. For instance, on human mRNA half-life prediction, Orthrus MLM is the only self-supervised model to match the supervised performance (Pearson’s R 0.71). These results indicate that a simple linear regression trained on Orthrus embeddings can replace the expensive task-specific training of neural networks for mRNA property prediction.

We observe improved linear probing results as we scale the number of trainable parameters by comparing Orthrus Small and Orthrus model variants (Figure 2). We find that this trend of improvement is strongest in MRL and GO Molecular Function predictions. We note that for other self-supervised models such as Hyena DNA [14] or Nucleotide Transformer [29], increasing the number of parameters does not consistently improve performance (Figure A3 & Table A1). However, we do observe an improvement in performance for Nucleotide Transformer when comparing their 2.5 billion parameter model pre-trained on 1000 genomes data versus multi-species. This is additional evidence that using evolutionary information can help improve model performance on mRNA property prediction tasks [44].

### Fine-tuning Orthrus for state-of-the-art mRNA Property Prediction

To assess whether the Orthrus pre-training objective provides utility beyond an effective representation, we evaluate its performance by fully fine-tuning it and comparing it to a supervised model with a matched architecture. We compare its performance against a published method for the mRNA half-life prediction, Saluki [7], and find that the fully fine-tuned Orthrus model outperforms Saluki on the mRNA half-life task (Figures 2 C, A5). Furthermore, we retrain the Saluki architecture, train an architecturally equivalent model to Orthrus for the other sequence property prediction tasks and identify that Orthrus has a significant performance advantage (Figures A2, A5).

To simulate downstream tasks for which there is a lack of experimental data, we perform finetuning on mRNA property prediction tasks where only a subset of the original training data set is available. We observe that *Ab initio* methods are ineffective in this regime, while Orthrus maintains competitive performance when trained on just 300 to 100 data points (Figure 2 C). The performance differences are even more stark when fine-tuning using just 30 training data samples, resulting in 71% of supervised performance on the human mRNA half-life dataset (Pearson’s R = 0.74 vs Pearson’s R = 0.53). These results demonstrate that Orthrus is effective in few-shot settings, offering strong performance on downstream tasks even when experimental data is scarce.

### Quantifying Orthrus Design Choices

We quantitatively evaluate the contributions of individual design choices in Orthrus by performing a comprehensive ablation study across three core properties: model training objective, data augmentations, and model architecture (Figure 3). To enable direct performance comparison across tasks with varying metrics and dynamic ranges, we normalize the results by converting them to Z-scores based on the distribution metrics from 10 independent runs and all model configurations (additional details in Section 7). This normalization allows us to compute a single aggregate Z-score for each configuration, providing a unified measure of overall model effectiveness. Our analysis of training objectives reveals that the contrastive learning (CL) objective alone yields higher overall performance when compared to a masked language modeling (MLM) objective with matched architecture and training data (0.9 vs 0.71). The MLM objective is statistically on par with CL objective on two tasks, while its performance significantly declines on RNA-centric benchmarks such as RNA half-life and mean ribosome load, underscoring the contrastive objective’s ability to capture signals relevant to mRNA metabolism. A combined CL and MLM objective further improves overall performance, suggesting the two approaches capture complementary information. The ablation over the augmentation set confirms the value of both splicing and orthology data sources. Specifically, including orthologous transcripts as positive pairs provides a substantial performance improvement over using only masking, highlighting the benefit of incorporating evolutionary signals (Z score -0.11 vs -0.55). Finally, a comparison of model architectures demonstrates a significant performance advantage of using a Mamba-backbone over parameter matched baselines, including a Saluki-like larger model and a dilated CNN (0.9, -0.23, -0.53). This ablation also confirms our earlier findings on parameter scaling, with the Mamba 10.1 million parameter architecture model consistently outperforming the Mamba Small 1.3 million variant (0.9 vs 0.72). For transparency and ease of comparison, the raw performance metrics for each experiment are provided in Figure 3 B.

**Fig. 3:**
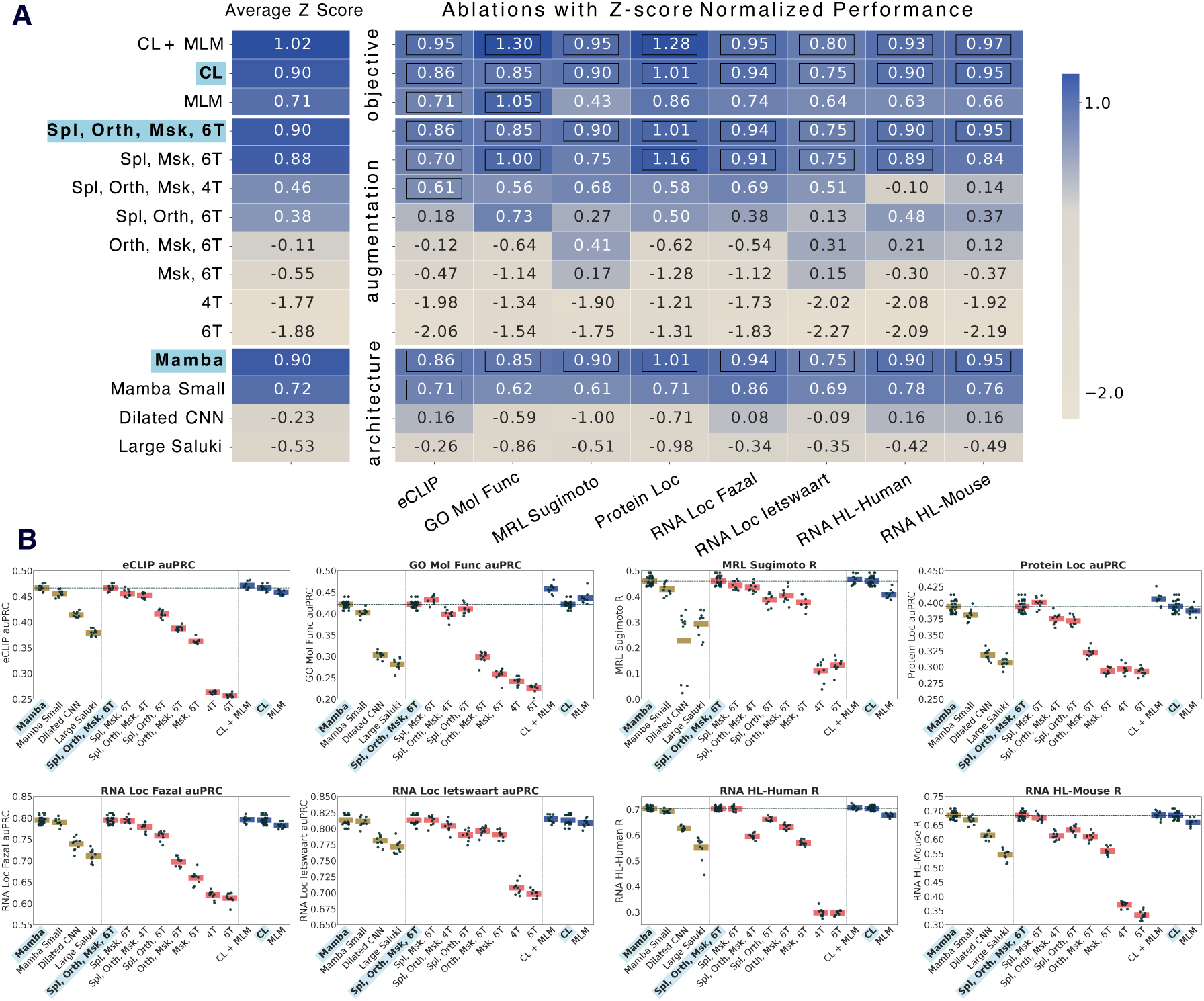
Ablation study of Orthrus pre-training and architecture. We perform an ablation over core properties of the model that result in effective performance: the pre-training objective function, the set of data augmentations, and the model architecture. Abbreviations: CL = contrastive learning, MLM = masked language modeling, Spl = splicing, Orth = orthology, Msk = masking, 6/4T = track, CNN = convolutional neural network. **(A)** To create a comparable aggregate score across tasks with different dynamic ranges, we transformed performance metrics: Area Under the Precision-Recall Curve (auPRC) for classification and Pearson’s R for regression into Z-scores. For each model configuration and downstream task, we performed linear probing with 10 different random seeds. The resulting distribution of scores for each task was used to normalize the values, creating a normal distribution of performance. The heatmap visualizes these Z-scores for each model (rows) across each task (columns), with the average Z-score shown on the left. Black boxes indicate that a model’s performance is not statistically significantly different from the reference model (highlighted in blue and bolded), which corresponds to the Orthrus contrastive learning variant, as determined by a onesided paired t-test where samples are paired by random seed. **(B)** Jitter plots showing the raw performance values for the models presented in Panel A on each of the downstream tasks. We use auPRC as the metric for classification tasks and Pearson’s R for regression tasks.

### Orthrus Latent Space Encodes Functional Similarities

Having shown that Orthrus captures properties of individual transcripts, we then investigated its ability to detect variation in function among different isoforms of the same gene [35]. To explore this, we compute Orthrus embedding similarities for each pair of protein-coding transcripts of the same gene (Figure 4 A). As a control, we compare these with the transcript pairs of random genes, expecting a lower similarity. We also hypothesized that transcript pairs from genes sharing the same GO terms would be more similar than random pairs, but less similar than most intragene pairs. Our analysis confirms significant differences across all pairwise comparisons of the three groups (p *<* 2.2e-16, two sided Mann-Whitney U test), indicating that the Orthrus training objective preserves within gene sequence diversity (Figure 4 B). Notably, we observe an overlap between intragene and intergene similarities, indicating that some alternatively spliced transcripts have distinct embeddings, RNA properties, or functional differences in protein products. As such, *within gene* diversity could potentially help delineate differential isoform protein functions, an active area of research.

**Fig. 4:**
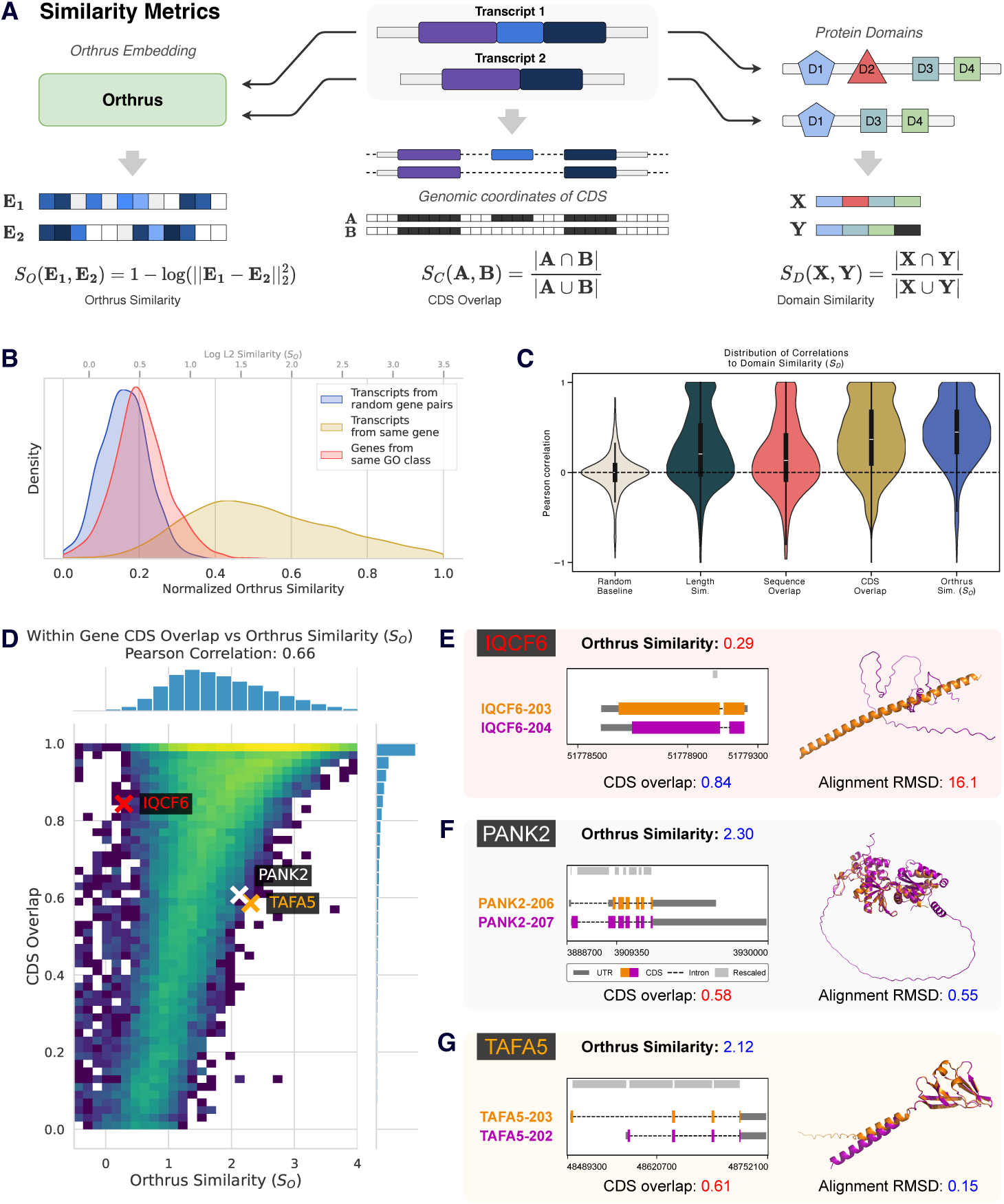
Analysis of functional similarities in Orthrus embeddings. **(A)** Methodology for comparing gene isoform similarities using Orthrus embeddings, protein domain annotations, and CDS overlap. Orthrus embeddings for transcripts within the same gene are compared using log of L2 distance, while protein domain similarity and CDS overlap is computed using the Jaccard index. **(B)** We visualize within gene similarities (yellow), between gene similarities (blue), and similarities of genes from the same GO class (red). **(C)** Visualization of Pearson’s R distributions correlating protein domain similarities with baseline or Orthrus embedding similarities for 1000 randomly sampled genes with multiple isoforms. **(D)** Heatmap illustrating the relationship between CDS overlap and Orthrus similarity in pairs of alternatively spliced transcripts. **(E-G)** Comparative analysis of transcript pairs where the CDS overlap and Orthrus similarity metrics are divergent. AlphaFold3-predicted structures for isoforms are superimposed. Orthrus similarity captures structural similarity as quantified by root-mean-squared deviation (RMSD). Genomic coordinate plots for individual isoforms are also shown.

To investigate whether Orthrus similarities for intragene transcripts reflect functional similarity, we compared annotated protein domains across each transcript pair (Figure 4 A). We find that transcripts with a high degree of protein domain overlap (*S_D_*) also have highly similar Orthrus embeddings (*S_O_*) with a median Spearman’s rho and Pearson’s R correlations of 0.37 and 0.45 respectively. This correlation is significantly higher than when compared to baseline metrics such as transcript length and overall sequence overlap, indicating that Orthrus better captures functional differences encoded by protein domains (Figure 4 C). We expect that transcript sequence overlap, especially in coding regions, is likely to contribute to similarities in RNA embeddings, and visualize the relationship between CDS overlap (see Methods) and Orthrus similarity for intragene pairs of protein-coding isoforms across all genes (Figure 4 D). An overall positive correlation is observed (Spearman’s rho = 0.68, Pearson’s R = 0.66), confirming that Orthrus identifies the coding sequence as a major determinant of function. However, we observe that Orthrus captures variability in functional similarity that extends beyond sequence similarity.

In the panels shown in Figure 4 (E-G), we investigate individual examples where transcript pairs have high CDS overlap but low Orthrus embedding similarity (*IQCF6*) and the reverse, where sequences share a smaller fraction of the sequence and yet their embeddings are close, as indicated by Orthrus similarity (*PANK2, TAFA5*). Isoforms in *IQCF6* have a very similar coding sequence (0.93) but a low Orthrus similarity (0.84, 1.52*^th^* percentile), suggesting our model predicts them to be functionally distinct. To evaluate the functional differences between these splice isoforms, we generate structure predictions of the transcripts’ coding sequences with AlphaFold3 [45] and structurally align them using PyMOL [46]. The structures demonstrate low overall alignment, as evidenced by a Root Mean Square Deviation (RMSD) of 16.1 angstroms (^°^A), inline with our Orthrus-based scoring. Meanwhile, isoforms in both *TAFA5* and *PANK2* exhibited low coding sequence overlap (Jaccard Index *<* 0.6, bottom 20*^th^* percentile). However, Orthrus embedding similarity is predicted to be high (60*^th^* and 55*^th^* percentiles respectively). Similarly, we visualize the predicted structures for the pairs of transcript coding sequences and find low overall RMSD values of 0.15^°^A and 0.55^°^A, confirming structural similarity despite low coding sequence overlap. Cumulatively, these findings suggest that Orthrus embeddings encode functionally relevant information.

### Orthrus Embeddings Capture Functional Isoform Diversity

We investigate whether Orthrus is able to capture well-studied examples of intragene divergent function by clustering learned embedding similarities. To illustrate this, we select two genes that are experimentally annotated with distinct isoform function and localization. We first examine *BCL2L1*, known for its alternatively spliced isoforms with distinct roles in the apoptosis pathway [47, 48].

The dominant isoforms encode an apoptosis-inhibiting protein, Bcl-X(L), while a minority encode a pro-apoptotic protein, Bcl-X(S). By clustering *BCL2L1* RNA isoforms using Orthrus embedding similarity, we identify two main functional groups: one containing *BCL2L1-202* and *BCL2L1-205*, distinct from the apoptosis-inhibiting transcripts cluster (Figure 5 A).

**Fig. 5:**
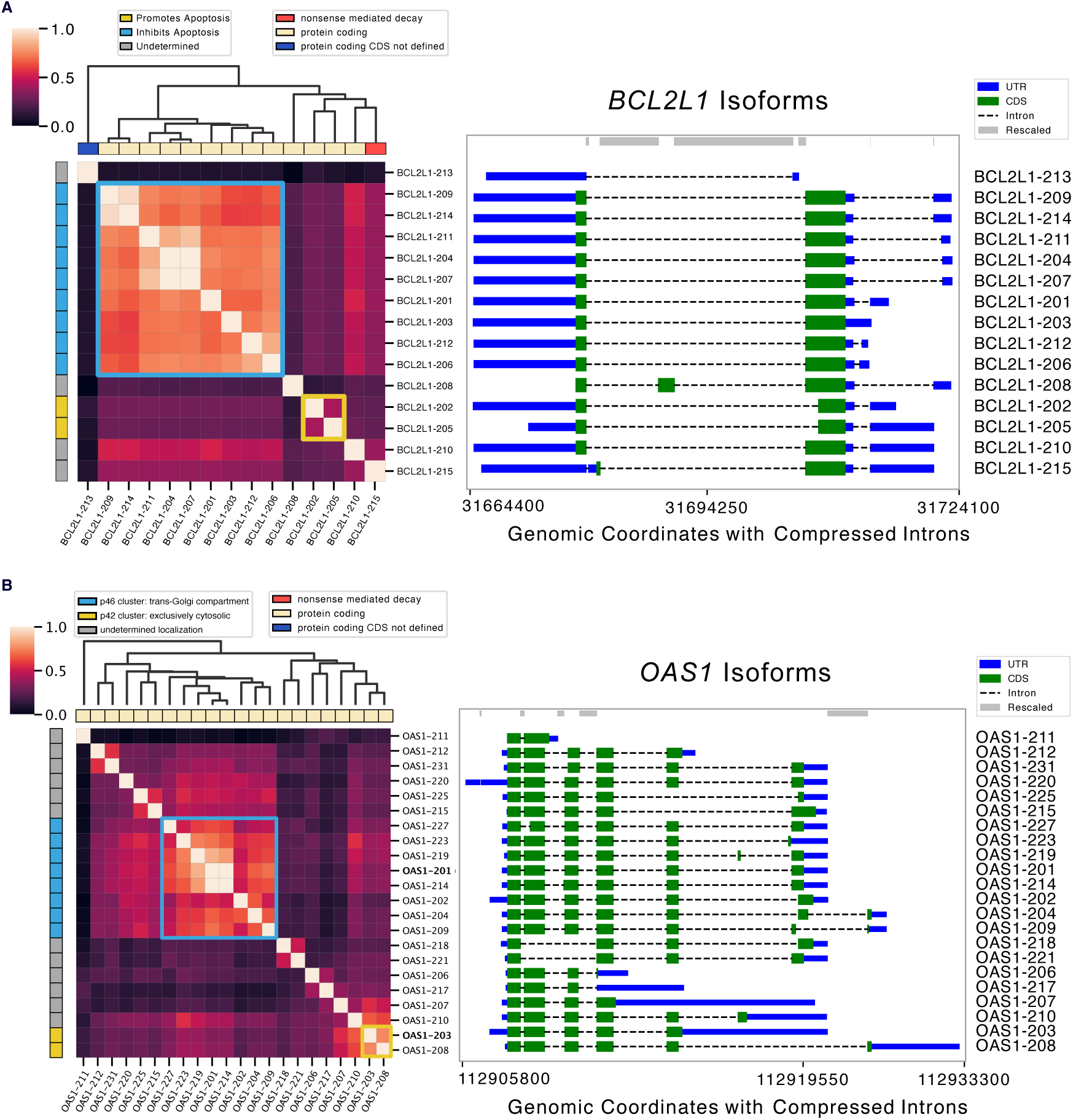
Functional clustering of splice isoforms using Orthrus. Heatmaps visualize normalized Orthrus embedding similarity between all splice isoforms within the analyzed gene. Transcript biotype is annotated along the x-axis, while functional annotations for isoforms are displayed along the y-axis. Isoform genomic coordinate plots are shown on right. The clustering matrix, derived from Orthrus embedding similarities, is represented by the dendrogram. Highlighted cells in the heatmap indicate clusters with divergent transcript functions. **(A)** *BCL2L1* isoforms where apoptosis-inhibiting isoforms cluster together, while noncoding and apoptosis-inducing isoforms display low similarity. **(B)** *OAS1* isoforms clusters represent distinct cellular localization trends associated with viral response.

As another example, the two major isoforms of *OAS1*, p42 and p46, have previously been shown to have distinct antiviral functions [49]. Prenylation, a post-translational modification that adds a hydrophobic lipid group to a protein, allows the p46 (*OAS-201*) isoform to anchor to membranous organelles such as the trans-Golgi compartment and block the replication of viruses such as SARSCoV. However, the p42 (*OAS-203*) isoform lacks prenylation and does not inhibit viral replication, highlighting functional divergence between isoforms [49]. Using Orthrus embedding similarities, we show that these two distinct isoforms form their own marked clusters (Figure 5 B). These case studies demonstrate the model’s ability to cluster isoforms by function. We extend the evaluation of this capability by predicting the results of a genome wide exon deletion screen on cellular fitness (Appendix A.1, Figure A1). These findings demonstrate that Orthrus embeddings may serve as a valuable resource for identifying functionally divergent isoforms, a critical area in alternative splicing research.

## Discussion

In this work, we introduce Orthrus, a mature RNA foundation model that is trained to capture the diversity of RNA through an evolutionary and functional lens [50, 51]. We create a self-supervised training objective that learns similarities between evolutionarily related sequences identified in the Zoonomia project [32]. In addition, we utilize alternatively spliced transcripts to learn sequences responsible for shared functions between splicing isoforms [52]. By training on sequences generated by evolutionary and alternative splicing processes, Orthrus utilizes stronger biologically motivated inductive biases compared to SSL reconstruction methods. This makes Orthrus less reliant on limited genetic sequence diversity during pre-training, and capable of learning strong representations without fine-tuning on experimental data.

Previous self-supervised works for genomic sequence property prediction have focused on reconstruction objectives like masked language modeling or next token prediction [12, 29]. However, most positions in the human genome are under little to no negative selection, and are not as informative for model training [25, 26].

By minimizing the embedding distance between mature RNAs related by speciation and alternative splicing, Orthrus produces representations that are highly predictive of properties like mRNA half-life and mean ribosome load, achieving state-of-the-art performance when fine-tuned (Figures 2 B and A5). We observe that pre-training is especially helpful in low data regimes when there are 300 or fewer data points with labels (Figure 2 C). We demonstrate that carefully designed selfsupervised pre-training mitigates the data efficiency challenges present in genomics, and that scaling to additional species can be an effective dataset expansion strategy.

A key question is why our proposed contrastive objective, which minimizes embedding distance between related transcripts, is effective for predicting functional properties. Our central hypothesis is that the contrastive framework identifies conserved functional segments by learning from the sequence variations introduced by alternative splicing and speciation. This aligns with recent theoretical work suggesting that contrastive methods excel at disentangling shared functional information from sequence-specific “style” [53]. Our own results provide strong evidence for this: we find that the learned similarity between isoforms correlates directly with their shared protein domains (Figure 4). Furthermore, our categorical Jacobian analysis of *TAF5* reveals that Orthrus learns the interdependencies within the functionally critical WD40 domains, particularly highlighting the fitness-promoting exon 8 as a coherent functional unit (Figure A2). By learning to encode these functional invariances, Orthrus can predict complex mRNA properties such as mean ribosome load and mRNA half-life (Figure 2).

A possible limitation of our approach is that by maximizing similarity in representation space between functionally related sequences, such as splice isoforms, we remove important signals for predicting properties. We investigate this by predicting the cellular fitness impact of exon skipping events from a genome-scale perturbation screen [54] (Figure A1). Orthrus successfully classifies exons as fitness-promoting based on the change in embeddings upon exon deletion, demonstrating it captures transcript-specific functional information (Figure A1). This is an important capability, as while for some mRNA processes such as mRNA half-life, previous works have found that isoform choice has little discernible effect [55], for others like neurological development it is functionally critical [56, 57].

Although the Orthrus training objective encourages clustering of alternatively spliced isoforms in the model’s latent space, our results show that Orthrus effectively captures transcript-specific functions. As illustrated in Figure 4, Orthrus representation similarities reveal functional divergence among isoforms, even when they share extensive sequence overlap. Further, Figure 5 demonstrates that Orthrus can group alternatively spliced transcripts based on their distinct functional roles. This capacity to discern transcript-level functionality is especially valuable given the scarcity of transcript-level functional annotations, as it enables more detailed biological insights. Conditioning these analyses on properties such as protein localization or mRNA stability, for instance, could guide investigations into even more nuanced aspects of transcript-level functional variation.

In this work, we propose a novel, self-supervised contrastive objective for learning mature RNA isoform representations. We show that this approach is an effective strategy to address two major challenges for cellular property prediction: data efficiency and model generalizability. We demonstrate that Orthrus representations are effective in the low data setting, paving the path to true few-shot learning for mRNA property prediction. Our ablation studies verify that the contrastive objective, evolutionary augmentations, and Mamba architecture are all key contributors to this performance (Figure 3). Finally, our best model, Orthrus MLM, outperforms much larger self-supervised models using over 700x fewer parameters than the second best model and surpasses even *Ab initio* supervised models on key benchmarks.

## Methods

Contrastive learning has been shown to be a bound on mutual information between two random variables X and Y corresponding to *I*(*X*; *Y*) = E*_p_*_(*x,y*)_ log *<u>^p^</u>*<u>^(*x,y*)^</u> . We utilize a variation of the classical InfoNCE loss, E log <u>^exp(*f*(*x*^</u>*<u>^i,yi^</u>*<u>^))^</u>, where a model *f* is tasked with classifying the correct *y_i_* which was jointly drawn with *x_i_* [58]. Herein, the observations *x_i_, y_i_* correspond to splice isoforms or orthologous sequences which are interpreted as functionally related while *f* is a neural network that we optimize to minimize the loss.

We propose to use four different augmentations and thoroughly investigate their impact on downstream tasks. They include: alternatively spliced transcripts across ten organisms, orthologous transcripts identified from the Zoonomia project including over 400 species, naive orthology informed by gene identity, and masking 30% of the input sequence (Figure 1) [32, 39].

In the following section we elaborate on dataset construction, model choice, contrastive learning objective, and downstream evaluations.

### Splicing and Orthology Contrastive Dataset

In the computer vision domain, contrastive learning strategies have had significant success by identifying augmentations that do not have a strong semantic effect, such as cropping, rotation, or Gaussian blur [31]. In this work, we use RNA splicing isoforms and orthologous transcripts as sources of functional similarity [32, 38, 39]. By sampling RNA isoform sequences produced by alternative splicing and speciation processes, we identify sequence variation that is likely to maintain core functional properties. In addition, we use naive orthology to pool RNA transcripts from evolutionarily related genes [52]. Here, for cases where gene names are consistent between species, we pool the transcripts generated by alternative splicing into the same transcript set. By minimizing the distance between functionally similar sequences, the model can learn regulatory regions critical for mRNA property and function prediction.

We generate a six-track mature RNA representation, consisting of four one-hot encoded tracks encoding genomic sequence, a track indicating the 5’ location of splice sites, and a track indicating the first nucleotide of every codon in the CDS. The addition of splice site and coding sequence locations has been shown to be beneficial for mRNA property prediction tasks [7].

To sample positive pairs from the orthology and splicing dataset, we first identify the set of all positive samples **Y_j_**for a reference transcript *x_j_*. Since Zoonomia derived orthologs lack UTRs, during sampling *y_j_*^Orthology^ we introduce a UTR combination transform augmentation to prevent the model from penalizing UTR importance. This transform creates a chimeric transcript by attaching the UTRs from a splice isoform *x_j_* to the coding sequence of its corresponding ortholog *y*^Orthology^. **Y_j_** can be variable in length since some transcripts will have a greater number of splice isoforms and orthologous sequences than others. During a forward model pass, we sample *y^k^* from **Y_j_** and use that as a positive pair for *x_j_*.

### Mamba Encoder

We pre-train a Mamba state space model, which has been demonstrated to be successful in applications with long context requirements [14, 33]. mRNA sequences can reach over 12,000 nucleotides in length, making application of the Transformer architecture challenging due to its quadratic scaling in memory with sequence length [59]. Mamba, an extension of state space model families and S4 [60], maps a sequence *x*(*t*) ∈ R to *y*(*t*) ∈ R using a latent state *h*(*t*) ∈ R*^N^* .

A fundamental trade-off in architecture choice for sequence modeling is avoiding compressing sequence context and compute requirements. Transformers are able to avoid compressing context, leading to better performance, but trade-off slower training and higher memory usage [33, 59]. Alternatively, S4 models define a sequence to sequence transformation parameterized by (**A**, **B**, **C**, Δ). The fundamental operation consists of iteratively updating the hidden state:

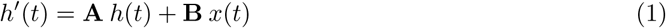

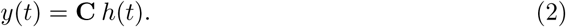

Δ is used to discretize the input for discrete domains such as natural language, or genomics. The Mamba architecture iterates on the S4 family of models by introducing selectivity over input by making *B*, *C*, and Δ a function of the input, resulting in

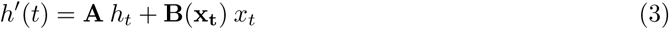

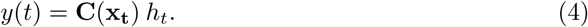

Allowing parameters to be input dependent introduces desirable modeling qualities for genomic domain: variable spacing, filtering context, and linear memory scaling with sequence length O(*n*). Variable spacing refers to Mamba’s ability to effectively perform on the selective copying task, where causal elements are arbitrarily spaced [33]. Binding motifs in genomic sequences can be spaced without a constant offset, requiring the model to be able to learn motif interactions with variable spacing [61]. The non-unformity of signal informativeness in genomic sequences requires models to be able to filter out irrelevant context [33]. Finally, the limited context, as opposed to Transformer models, allows the Mamba architecture to scale required memory linearly with increased input length [33, 59].

### Self-Supervised Pretext Training Objectives

**DCL:** for contrastive learning we use the decoupled contrastive learning (DCL) loss as it has been shown to require smaller batch sizes, is less sensitive to hyperparameters such as learning rate, and the positive loss term can be weighted by sample difficulty [42]. DCL iterates on the normalized temperature-scaled cross-entropy loss by splitting the contrastive objective into two terms: a similarity loss (positive) and a dissimilarity loss (negative) [62]. More formally, the positive and negative losses for sample *i* are calculated:

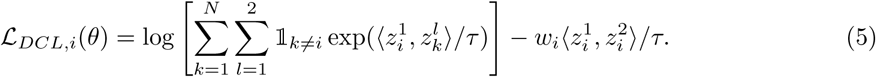

In the above *z*^1^ and *z*^2^ correspond to two embeddings of related sequences, *z_k_* are embeddings from unrelated RNA sequences, *τ* is the temperature parameter set to 0.1, and]_*_k_*_=*i*_ is an indicator function that evaluates to 1 when *k* ≠ *i*. The above loss is computed for all the samples in the batch for both the sampled views *l* ∈ 1, 2. *w_i_* is be used to weight the importance of individual augmentations. In practice it is set to one besides Orthology augmentations, for which we empirically find 0.8 results in the best downstream performance. *N* corresponds to all the negative samples in batch, thus maximizing batch size during contrastive learning typically leads to improved performance.

Normalized projections *z_i_* are outputs from the MLP projector *g_θ_*and are used to compute the contrastive loss, utilizing samples from the rest of the batch as negative examples:

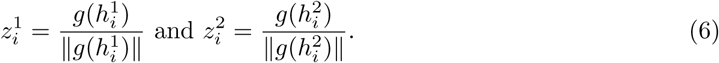

For downstream RNA property evaluations, the projector *g_θ_* is discarded and outputs from *f_θ_* are used instead. This practice is consistent with prior literature [31].

### Joint MLM and contrastive learning

To capture both local nucleotide-level patterns and global transcript-level features, we train Orthrus using an objective that combines Masked Language Modeling (MLM) and Contrastive Learning (CL). Unlike autoregressive objectives such as next-token prediction, which requires causal masking, combining MLM with CL allows the model to learn from the full sequence context. Both the MLM and CL learning signals are derived from a shared Mamba sequence encoder.

The MLM task trains the model to reconstruct the original nucleotide identities at randomly masked positions within an input sequence. For a given input transcript, we randomly select 15% of its nucleotide positions for masking. The one-hot encoded vectors at these positions are replaced with a zero vector, creating a corrupted input sequence *X*. The model’s sequence head then predicts the probability distribution over the four nucleotides for each position in the sequence. The MLM loss is calculated using a standard cross-entropy function, but only over the set of masked positions, M.

Let *X_i_* be the original one-hot encoded nucleotide vector at a masked position *i* ∈ M, and let *P*^^^*_i_* be the model’s predicted probability vector for that same position from *X*. The MLM loss is formulated as:

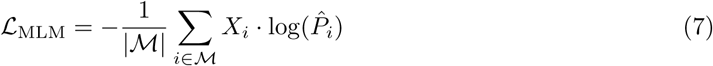

The final training loss is a weighted sum of the two objectives. Because the raw loss values for CL and MLM often have different numerical scales, we introduce a weighting coefficient *α* to balance their contributions and ensure stable training. The total loss is defined as L_total_ = (1 − *α*) · L_CL_ + *α* · L_MLM_. Based on empirical analysis of the loss norms, we set *α* = 0.95 to apply a higher weight to the MLM objective.

### Downstream Evaluation Tasks

**mRNA half-life (mRNA HL)** is an important cellular property to measure due to its implications for protein expression regulation. Recently, it has been shown that the choice of experimental methodology for measuring mRNA half-life can have an outsized impact [7]. To address this challenge, Agarwal and Kelley (2022) utilized the first principal component of over 40 different mRNA half-life experiments. The dataset consists of 10,432 human and 11,008 mouse mRNA sequences with corresponding measurements. The low data availability and high inter-experiment variation underscore the importance of data efficiency, and generalizability in computational models to be developed for this task.

**Mean ribosome load (MRL)** is a measure of the translational efficiency of a given mRNA molecule. It measures the number of ribosomes translating a single mRNA molecule at a point in time. Accurate MRL measurement is crucial as it offers insights into the efficiency of protein translation, a key process in cellular function. The dataset in question, derived from the HP5 workflow, captures this metric across 12,459 mRNA isoforms from 7,815 genes [63]. This dataset was derived from a single experiment, so we can expect a higher amount of noise associated than the mRNA half-life dataset.

**Protein localization** describes a protein’s subcellular location, which can be determined using cells that are immunofluorescently stained. Protein function is often linked with its localization, underscoring the importance of this task. We downloaded a dataset of 10,409 genes, whose protein localization was determined by the Human Protein Atlas [64]. We included the 12 most common locations including Nucleoplasm, Cytosol, Vesicles, Mitochondria, Plasma Membrane, Golgi apparatus and others. We utilized one transcript per gene (defined to be the canonical isoform by the Appris database [65]).

**Gene ontology (GO)** terms are a hierarchical classification system used for assigning function to genes and their products [66–68]. In this work, we utilize GO classes to visualize model latent embeddings and classification. GO term hierarchical systems allow for fine-grained annotation of function, with broader terms at the top of the hierarchy and increased specificity closer to the bottom. To annotate genes with gene ontology terms, we subset GO classes three levels from the root, labeling all available genes. GO categories are assigned to individual genes, so to select an isoform for each gene we use the APPRIS database for prioritization [69] .

**eCLIP Binding (eCLIP)** curated by the authors of mRNA-Bench [70], is derived from ENCODE [71]. The enhanced UV crosslinking and immunoprecipitation (eCLIP) protocol identifies the binding sites of RNA-binding proteins (RBPs) on a transcriptome-wide scale [72]. RBPs are crucial regulators of mRNA processing, controlling functions like alternative splicing, stability, and nuclear export. The benchmark uses data for 168 RBPs across two cell lines, K562 and HepG2. For the prediction task, the top 20 RBPs per cell line were selected based on the number of binding events, the data from both cell lines was aggregated, and the task was framed as a binary classification of whether a given RBP binds to a specific transcript.

**mRNA Subcellular Localization Ietswaart et al.** An mRNA molecule’s path from transcription to translation involves traversing multiple cellular compartments, including chromatin, the nucleus, and the cytoplasm. This dataset, processed by mRNA-Bench from Ietswaart et al., uses direct RNA sequencing to create an isoform-resolved map of this process [70, 73]. The assay captures RNA flow dynamics by measuring the rates at which transcripts are released from chromatin, exported from the nucleus, and loaded onto polysomes for translation. While a transcript passes through all of these locations, its steady-state level leads to enrichment in a primary compartment. The task is to predict this subcellular enrichment for a given mRNA transcript.

**mRNA Subcellular Localization, Fazal et al.** assembled by the authors of mRNA-Bench [70], leverages APEX-seq, an engineered peroxidase that performs proximity-based biotinylation to map the RNA landscape with high spatial resolution [74]. The assay provides a quantitative atlas of transcript localization across eight distinct subcellular compartments, including the nuclear lamina, nuclear pore, and outer mitochondrial membrane. For the purposes of evaluation, the proportional abundance data was converted into a multi-label classification task to predict the presence of a transcript in each of the eight compartments.

### Associating Orthrus RNA Embeddings with Transcript Similarity and Protein Domains

To evaluate how well Orthrus RNA embeddings capture functional diversity among transcript isoforms, we analyzed the similarity of transcript pairs within and between protein-coding genes, excluding homologous genes when comparing random gene pairs or genes sharing the same GO term. The test dataset for this analysis was prepared as follows:

1. **Intragene Pairs**: We sampled 1,000 genes to obtain pairs of protein-coding transcripts.
2. **Intergene Pairs**: We randomly sampled 1,000 pairs of non-homologous genes, selecting the MANE transcript for each gene, which represents the most likely relevant isoform.
3. **Intergene Pairs**: We sampled 5,000 GO terms, each containing 10 to 1,000 genes, and selected five non-homologous gene pairs per term.

For each transcript, we computed Orthrus embeddings and calculated pairwise distances between embeddings using the L2 norm. We calculated a similarity score for each transcript pair as 1 − log(L2 distance). This ensures more interpretable results, where higher similarity scores correspond to closer RNA embeddings in the latent space, allowing us to compare the three groups of transcript pairs.

To assess whether similarities in Orthrus embedding reflected shared functional features, we annotated each transcript with protein domain information using Ensembl data and the Pybiomart package. We used the Jaccard Index to quantify the similarity of protein domain presence or absence between each pair of transcripts within a gene. The Jaccard Index is defined as the size of the intersection divided by the size of the union of the protein domain sets present in each transcript pair:

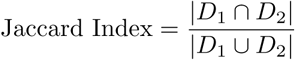

where *D*_1_ and *D*_2_ are the sets of protein domains present in each transcript. Higher values indicate greater similarity in protein domain composition. We calculated this metric using “intragene pairs” and “intergene pairs” to further study how protein domain composition correlated with embedding similarity. We analyzed the Pearson correlation between Jaccard indices and embedding similarities separately for intragene and intergene pairs to determine if transcript pairs within the same gene exhibited higher concordance. To calculate sequence overlap, we take either the full set of transcript genomic intervals (sequence overlap) or just the coding sequence (CDS overlap) and calculate similarity using Jaccard index as indicated above. This way, two fully overlapping sequences will have a Jaccard index of 1 and non-overlapping will be 0.

To generate structural alignments we use the AlphaFold3 server to fold protein sequences as identified in Ensembl v113 [75, 76]. We then take the 0*^th^* seed from generated structures and generate an alignment using PyMOL [46].

To further explore the utility of Orthrus embeddings, we conducted a detailed analysis of the *BCL2L1* and *OAS1* genes [47, 49]. Transcripts from this gene were clustered based on their Orthrus embedding similarity scores, with clusters visualized and annotated according to transcript type and known functional roles.

### Ablation Study Z score Methodology

We perform a comprehensive ablation study to isolate the contributions of model architecture, data augmentations, and training objective across eight mRNA property prediction tasks. To enable robust comparisons across tasks with different metrics and performance ranges, we normalize all results using a Z-score transformation. This procedure quantifies each model’s performance relative to a baseline distribution established from a reference set of all 13 model configurations shown in Figure 3. For each task, this reference distribution is created by pooling the performance scores from ten independent linear probing runs for every model in the reference set. Performance is measured using the Area Under the Precision-Recall Curve (auPRC) for classification tasks and Pearson’s R for regression tasks. For regression metrics, we first apply the Fisher Z-transformation (np.arctanh) to the Pearson’s R values. This standard statistical procedure stabilizes the variance and converts the skewed sampling distribution of correlation coefficients into an approximately normal distribution [77]. Using the pooled reference distribution for a given task, we calculate its mean *µ* and standard deviation *σ*. The performance score *x* from each of the ten runs for every model is then converted to a Z-score using the formula: *Z* = *<u>^x^</u>*<u>^−*µ*^</u> . The resulting Z-score measures a model’s performance in standard deviations from the reference mean, allowing for direct and meaningful comparison across all tasks. A final aggregate Z-score is then computed for each model configuration by averaging its task-specific Z-scores.

To assess statistical significance, we compare each model against a top-performing reference model using a one-sided Welch’s t-test on their respective distributions of 10 scores. For the results shown in Figure 3, the reference model is the main Orthrus variant (denoted as CL, Spl Orth Msk 6T, or Mamba Large), highlighted in bold. A resulting p-value greater than 0.05 indicates that a model’s performance is not statistically significantly different from the reference, placing it in the top performance tier for that task. These instances of non-significant difference are marked with black boxes in Figure 3A.

## Data Availability

For constructing the pre-training data, a species FASTA file and a corresponding genePred file were required to specify genomic coordinates. For the 10 reference species containing splicing information, data was downloaded from the UCSC Table Browser (https://genome.ucsc.edu/cgi-bin/hgTables). Gencode files were downloaded for human (v43 for hg38) and mouse (vm23 for mm10). For other species, RefSeq files and corresponding genome assemblies were downloaded, including:

- Chicken: galGal6 (transcriptome updated on 2020-04-01)
- *C. elegans*: ce11 (transcriptome updated on 2020-07-02)
- Chimpanzee: panTr06 (transcriptome updated on 2020-04-01)
- Cow: bosTau9 (transcriptome updated on 2020-04-01)
- Dog: canFam4 (transcriptome updated on 2022-01-26)
- Fruit fly: dm6 (transcriptome updated on 2021-02-11)
- Rat: rn6 (transcriptome updated on 2020-04-01)
- Zebrafish: danRer11 (transcriptome updated on 2020-04-01)

Homology information was obtained from Ensembl BioMart (http://www.ensembl.org/biomart/ martview) to identify human gene orthologs and corresponding species.

Zoonomia files were downloaded from the web portal provided through the TOGA publication. For mouse and human mappings, a reference file was obtained from the overview table (https://genome.senckenberg.de/download/TOGA/human hg38 reference/overview.table.tsv). Individual BED files were downloaded via a path pattern: https://genome.senckenberg.de/download/TOGA/mouse mm10 reference/ followed by order/species/geneAnnotation.bed.gz, and were converted to genePred using a conversion script. FASTA files were obtained from the NCBI Genome Database (https://www.ncbi.nlm.nih.gov/ datasets/genome/) under the IDs specified in the overview table. Several assemblies were obtained from DNA Zoo (https://www.dnazoo.org/). Individual transcript sequences were serialized as NumPy arrays using https://github.com/bowang-lab/Orthrus/blob/main/orthrus/data.py#L884.

All linear probing and fine-tuning data were deposited at Zenodo (https://zenodo.org/records/ 13910050). RNA half-life data was collected from the Saluki repository (https://zenodo.org/records/ 6326409). Mean ribosome load data from Sugimoto *et al.* (2022) (https://www.nature.com/articles/s41594-022-00819-2) is available in the supplementary data (*Source Data Fig. 1*). Gene Ontology data was downloaded from http://current.geneontology.org/ontology/index.html, and parsed to identify the third level from the root node. Human protein localization data was downloaded from the Human Protein Atlas (https://www.proteinatlas.org/humanproteome/subcellular/data#locations). For the eCLIP and RNA localization datasets, please refer to the paper from which this data was obtained [70]. For transcript prioritization, the APPRIS database was used (https://appris.bioinfo.cnio.es/#/downloads).

## Code Availability

The code for pre-training, linear probing, and fine-tuning is publicly available at: https://github.com/bowang-lab/Orthrus under the MIT License. Additional resources, including embeddings for the human transcriptome, the APPRIS principality dataset, and the linear probing datasets, are available at the Zenodo repository: https://zenodo.org/records/14708163. The pre-trained Orthrus models are also hosted on Hugging Face at https://huggingface.co/antichronology/orthrus, with individual models available for inference at: https://huggingface.co/quietflamingo/orthrus-base-4-track and https://huggingface.co/quietflamingo/orthrus-large-6-track.

## Author Information

These authors contributed equally: Philip Fradkin, Ruian Shi, Taykhoom Dalal

## Authors and Affiliations

Vector Institute, Ontario, Canada

Philip Fradkin, Ruian Shi, Brendan J. Frey, Leo J. Lee, Bo Wang

Computer Science, University of Toronto, Ontario, Canada

Philip Fradkin, Ruian Shi, Brendan J. Frey, Bo Wang

Computational and Systems Biology Program, Sloan Kettering Institute, New York, United States

Ruian Shi, Taykhoom Dalal, Quaid Morris

New York Genome Center, New York, United States

Keren Isaev

Systems Biology, Columbia University, New York, United States

Keren Isaev

Electrical and Computer Engineering, University of Toronto, Ontario, Canada

Brendan J. Frey, Leo J. Lee

Peter Munk Cardiac Center, University Health Network, Ontario, Canada

Bo Wang

## Contributions

P.F. and R.S developed the concept of the work and the initial prototype. P.F. and R.S. contributed to design and implementation of the algorithm, construction of the pre-training dataset, and the contrastive learning objective. R.S and T.D. contributed to the objective design using masked language modelling and benchmarking using linear probing. The fine-tuning experiments were executed by P.F. and T.D. K.I. and P.F. carried out the functional transcript diversity experiments. P.F, R.S., and K.I. wrote the manuscript with input from all authors. B.W., L.J.L., Q.M. and B.F. supervised the study.

## Corresponding authors

Correspondence to Bo Wang, Leo J. Lee, Quaid Morris

## Acknowledgements

Resources used in preparing this research were provided, in part, by the Province of Ontario, the Government of Canada, through the Canadian Institute for Advanced Research (CIFAR) and companies sponsoring the Vector Institute. P.F. is supported by a Natural Sciences and Engineering Research Council of Canada (NSERC) Postgraduate Scholarship (Doctoral Program), R.S. is supported by the Ontario Graduate Scholarship, P.F. and R.S. are supported by a Vector Institute research grant.

T.D. is supported by a National Science Foundation Graduate Research Fellowship under Grant No. 2139291. B.W. is supported by NSERC (grants: RGPIN-2020-06189 and DGECR-2020-00294), the Peter Munk Cardiac Centre AI Fund at the University Health Network and the CIFAR AI Chair Program. Research was supported by an NIH/NHGRI Grant to Q.M. (R01 HG013328). Q.M. was partially supported by a NIH/NCI Cancer Center Support Grant (P30 CA008748). L.J.L. and B.F. are supported by the NSERC Discovery Grant. B.F. is a Canada Research Chair.

We thank C. Harrigan, A. Jung and A. Moses for helpful discussions. We thank A. Young, H. Maan and V. Chu for feedback on the manuscript. This work used resources from the High Performance Computing group at Memorial Sloan Kettering Cancer Center. We are grateful for the Vector Institute and the Memorial Sloan Kettering Cancer Center for providing the computing resources for this project.

## Appendix A

### A.1 Splice Isoform Fitness Evaluation with Orthrus

**Fig. A1:**
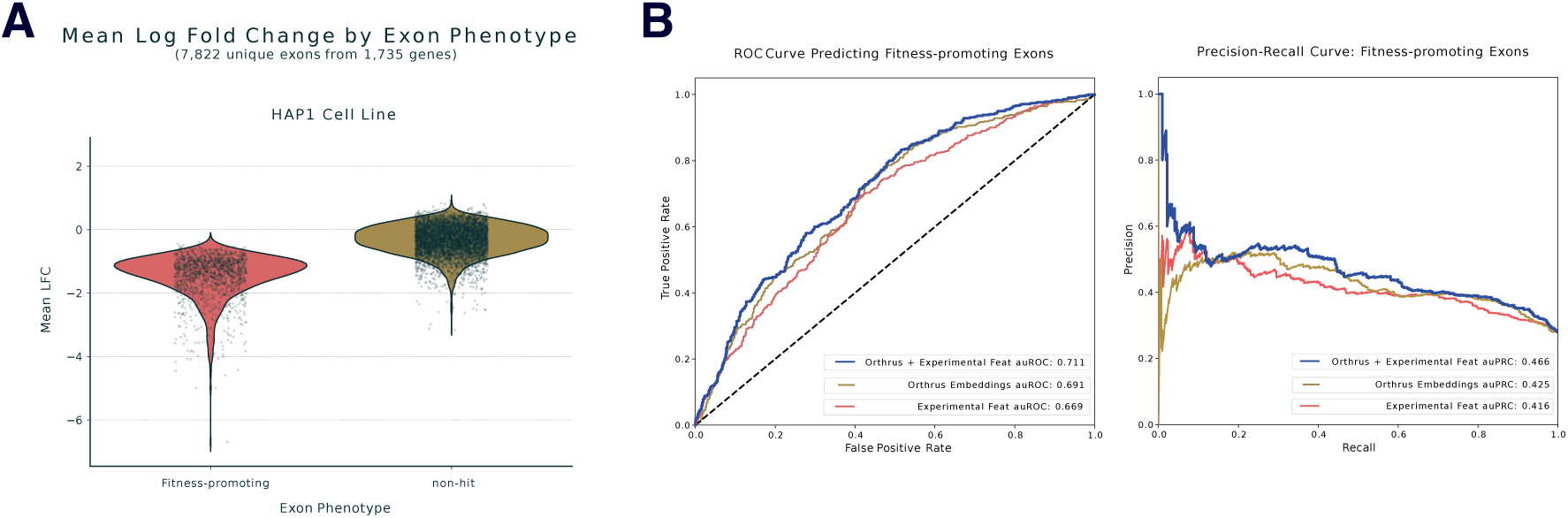
Orthrus embeddings are predictive of exon fitness phenotypes. **(A)** Distribution of mean log-fold change (LFC) for exons targeted in genome-scale CRISPR-based deletion screens in HAP1 and RPE1 cell lines. In an MPRA by Xiao et al. (2024), cells with deleted fitness-promoting exons are depleted, resulting in a significant negative LFC compared to non-hit exons. This distribution defines the ground-truth phenotypes for the classification task. **(B)** Performance of a random forest classifier trained to predict whether an exon is fitness-promoting. The model uses PCA-reduced embeddings from Orthrus for both the wild-type transcript and a version with the exon computationally skipped. Performance is evaluated using a Receiver Operating Characteristic (ROC) curve and a Precision-Recall curve. Orthrus based model is benchmarked against a model trained on curated experimental features (including exon percent spliced in values (PSI), gene expression, conservation, and splice site strength) and a model combining both embeddings and features.

To assess the ability of Orthrus to evaluate transcript function, we tested its performance on a genome-scale exon perturbation screen from Xiao *et al.* [54]. The screen utilized the ”CHyMErA” genetic perturbation platform, which employs Cas9 and Cas12a nucleases to perform targeted exon deletions. This experimental screen targeted 7,822 unique exons across 1,735 genes, identifying over 2,000 frame-preserving exons as ”fitness-promoting” (Figure A1 A). Deletion of these fitnesspromoting exons was shown to result in decreased cell viability. We replicated this experiment in-silico by generating paired Orthrus embeddings for each wild-type transcript *z_wt_* and its corresponding exon-deleted mutant *z_mut_*. The 7,822 sample pairs were split into 80% training and 20% testing sets, with stratification by gene to prevent information leakage between the splits. We used Principal Component Analysis (PCA), fit on the training embeddings, to project the wild-type and mutant embeddings into a lower-dimensional space. A random forest classifier was then trained on the projected embeddings to classify exons as either fitness-promoting or non-hit. The model trained on Orthrus embeddings alone achieved a predictive auROC of 0.69 and an auPRC of 0.425 (Figure A1 B). This level of performance is particularly notable given that the underlying experimental screen has a high reported false-negative rate of approximately 49%, which inherently limits the maximum achievable score by any predictive model. This performance surpassed a baseline model trained on curated experimental and sequence-based features, which achieved an auROC of 0.67 and an auPRC of 0.416 (Figure A1 B). The baseline feature set included Percent Spliced-In (PSI) values, gene expression levels, conservation scores, and splice site strength. Combining Orthrus embeddings with the experimental features further boosted performance, suggesting the two feature sets capture complementary predictive information (Figure A1 B). These results demonstrate that Orthrus can predict the functional impact of exon-level variations on cellular fitness, providing a high-throughput approach for evaluating splice isoform function.

To further validate what Orthrus learns, we perform a categorical Jacobian analysis on *TAF5* (A.2). We selected *TAF5*, a core component of the TFIID general transcription initiation factor, as the target gene for this analysis. This gene was thoroughly validated in the exon perturbation screen by Xiao et al., providing a robust experimental reference for functionally critical regions [54]. The perturbation screens highlighted the outsized importance of the WD40 repeat domains in *TAF5*, which were significantly enriched in fitness-promoting exons. Specifically, *TAF5* alternative exon 8 was identified as a key fitness-promoting exon that directly overlaps with one of these critical WD40 domains. Although considered alternatively spliced, exon 8 shows a high inclusion rate, with a Percent Spliced-In (PSI) value averaging 0.93 in humans. Despite its high inclusion, the skipping of exon 8 has profound functional consequences. Affinity purification-mass spectrometry experiments revealed that the resulting isoform, *TAF5-*Δ*E8*, fails to associate with any other TFIID component. This assembly failure directly impacts global gene expression, as *TAF5-*Δ*E8* is unable to rescue the majority of expression changes caused by the depletion of endogenous *TAF5*.

**Fig. A2:**
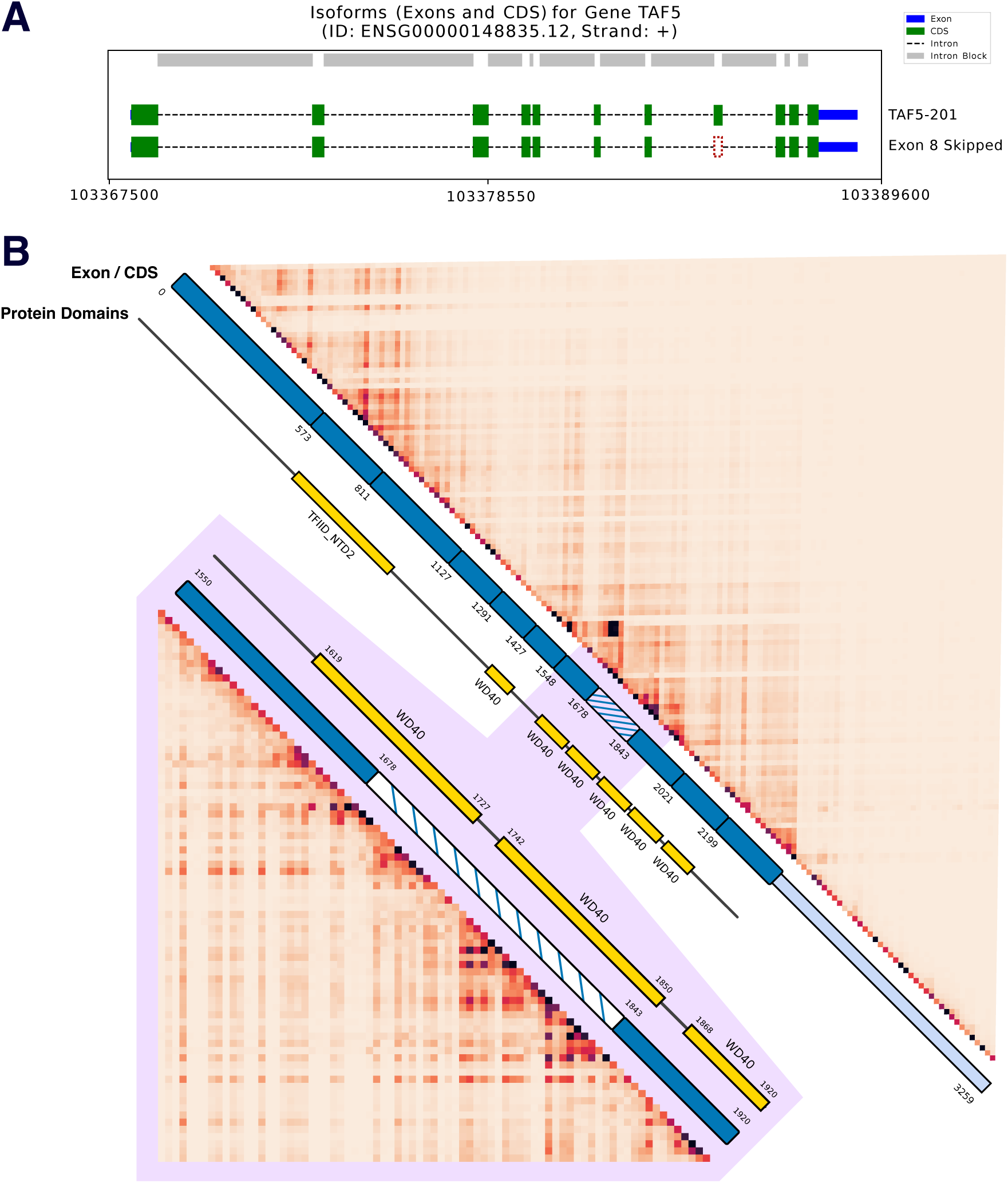
Orthrus captures functional intra-gene domains. **(A)** Genomic coordinates of TAF5 with exon 8 deleted, resulting in cellular loss of fitness as investigated by Xiao et al. (2024). **(B)** Categorical Jacobian analysis of the TAF5 transcript, a key fitness-promoting gene identified by Xiao et al. The heatmap reveals co-dependence between nucleotide positions, with strong signals corresponding to the boundaries of the WD40 repeat domains. This analysis demonstrates that the model captures the structure of functionally critical regions, such as exon 8 (highlighted), which was shown to be essential for TFIID complex assembly and overall cell fitness.

The categorical Jacobian analysis reveals the model’s learned dependencies between nucleotide positions by systematically mutating each site in the *TAF5* transcript and measuring the impact on the model’s output probabilities (Figure A2). Our analysis focuses on the region spanning the WD40 repeat domains, particularly the fitness-promoting alternative exon 8, whose inclusion is essential for TFIID complex assembly and global transcription regulation. Regions corresponding to the WD40 domains exhibit strong, localized interdependency patterns, indicating that Orthrus has learned to recognize these sequences as coherent functional units, entirely from unsupervised sequence data.

### A.2 Categorical Jacobian Calculation

To investigate the internal representations of Orthrus and quantify the interdependencies between nucleotide positions, we compute a categorical Jacobian matrix. This method, adapted from recent work in protein and DNA language model interpretability [78, 79], produces an *L* × *L* matrix for a sequence of length *L*, where the value at position (*i, j*) represents the maximum influence a single nucleotide substitution at query position *i* has on the model’s predictions at target position *j*. The calculation involves three main stages: *in silico* mutagenesis, calculating log-odds disruption, and aggregation.

**In Silico Mutagenesis and Prediction:** The initial step involves generating a comprehensive set of predictions for the wild-type sequence and all possible single-nucleotide variants.

For a given one-hot encoded wild-type (WT) sequence *X_wt_* ∈ {0, 1}*^L^*^×4^, a single forward pass is performed through the model. This yields a matrix of output probabilities *P_wt_* ∈ [0, 1]*^L^*^×4^, where index (*j, k*) of *P_wt_* is the model’s predicted probability of observing nucleotide *k* ∈ {A, C, G, T} at position *j*.

We systematically generate every possible single-nucleotide substitution. For each position *i* ∈ {1*, . . ., L*}, we create three mutant sequences by changing the original nucleotide to each of the three alternatives. This results in a total of 3×*L* mutant sequences. A forward pass is performed for each of the 3 × *L* mutant sequences to generate a corresponding output probability matrix, *P_mut_* ∈ [0, 1]*^L^*^×4^.

**Calculating Log-Odds Disruption:** To quantify the change in the model’s predictions, we calculate the difference in the log-odds (logit) space, which better captures shifts in model confidence than raw probabilities. The log-odds of a probability *p*is defined as:

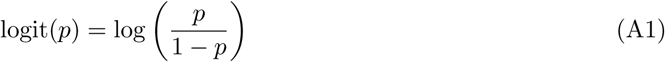

For each mutation (e.g., substituting the nucleotide at query position *i* to *k*_alt_) and for every target position *j*, we compute the change in log-odds for each of the four possible nucleotides *k*. This creates a log-odds difference matrix Δlogit ∈ R*^L^*^×4^ for each of the 3*L* mutations:

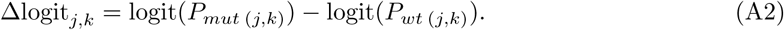

**Aggregation to the Final Dependency Matrix:** The final step involves aggregating these disruption scores into a single *L* × *L* matrix. To obtain a single scalar value representing the total disruption at a target position *j* caused by a specific mutation at *i*, we take the maximum absolute value across the four nucleotide predictions at position *j*. This score, captures the largest change in prediction at the target site, regardless of which nucleotide was affected. This calculation results in a disruption vector of length *L* for each of the 3*L* initial mutations.

To create the final matrix entry *M_i,j_*, we aggregate the disruption scores from the three possible substitutions at the query position *i*. We take the maximum value from the three corresponding disruption vectors. This ensures that *M_i,j_* represents the maximum potential influence that any mutation at position *i* can exert on the predictions at position *j*.

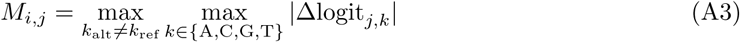

The final output is the categorical Jacobian matrix *M* ∈ R*^L^*^×*L*^, which can be visualized as a heatmap to reveal patterns of short and long-range nucleotide dependencies learned by the model.

### A.3 Model Architecture and Pre-training Hyperparameters

Here, we report the model architecture and hyperparameters used to train the Orthrus model. The Orthrus Small model uses a 3-layer Mamba block with a hidden dimension of 256, while the Orthrus and Orthrus MLM model use a 6-layer Mamba block with a hidden dimension of 512. A Mamba block consists of a selective state space model (SSM) core followed by a gated multi-layer perceptron (MLP). Both components are integrated into a residual architecture that applies layer normalization after the skip connection is added. All other hyperparameters are kept to their default value. Due to computational constraints, larger models were not evaluated. Orthrus uses a three layer projection head with a hidden layer size of 512, and projection dimension of 128. The network uses ReLU activations and BatchNorm between layers.

Orthrus was trained using either four A100 or H100 GPUs using PyTorch’s DDP. Orthrus Small models were trained using A40 and L40 GPUs. Due to the skewed distribution of transcript lengths, batches were constructed to minimize sequence padding. During each training iteration (corresponding to four batches, each of which is sent to an individual GPU), we create one smaller batch to contain longer transcripts, while the remaining three batches are larger with short transcripts. We found that using a sequence length threshold of 3,900 maximized GPU utilization, with resulting batch sizes of 50 and 150 respectively. Exact batch sizes were adjusted to maximize memory utilization on the GPU type used for each pre-training run. Orthrus was trained using mixed-precision.

All Orthrus models are pre-trained using the Adam optimizer with default hyperparameters for 20,000 steps. Model weights were checkpointed every 2,000 steps, and we select the final checkpoint based on validation contrastive loss. We note that the validation loss tended to plateau near the maximum step limit.

The model learning rate was selected from a grid of [1e-3, 1e-4, 1e-5] using validation loss, with a final value of 1e-3 used. We used a linear warm up scheduler with a factor of 0.1 for 2000 steps, and then cosine annealed learning rate for the remaining steps. We found weight decay to be important for constructing a robust latent representation, and a value of 5e-5 and 1e-5 was used for Orthrus and Orthrus Small respectively. The projection head uses a weight decay value of 1e-4. Gradient clipping was performed with a norm of five.

### A.4 Linear Probe Experimental Details

In this section, we describe the experimental procedure to evaluate linear probing results.

We first performed a 70-15-15 data split on datasets. The data sequences are then embedded by the various self-supervised learning (SSL) models. For Orthrus, we simply take the mean of the embeddings across the seqeunce dimension. For HyenaDNA, we take the mean and max of the embedding sequence dimension, as well as the last hidden state in the output sequence. For Evo2 models, following recommendations from the paper, we take both an intermediate layer and the final layers of the model for performing linear probing. Generating Evo2 embeddings requires H100 GPUs [19]. For supervised models such as Enformer [80] and Borzoi [81], we embed the RNA sequences directly, although the input typically consists of DNA which can explain the relatively low performance. This design choice is made to ensure that the models have access to equal input information and allows the models to represent alternatively spliced isoforms different embeddings. We pad the sequences with N nucleotides and perform length-aware pooling where we aggregate over only the regions that aren’t masked out. We also note here that we used the Flashzoi implementation of the Borzoi model from [82], which has been shown to replicate the results of Borzoi but with the added benefit of using FlashAttention2, enabling much faster evaluation. Furthermore, similar to the original Borzoi paper, we take the mean of the output from four replicates of the trained Flashzoi model as the embedding for each sequence. Other SSL methods could not handle input sequences of more than 500 or 1000 nucleotides. Thus, when input sequences exceeded the allowable context window, each sequence was chunked to the maximum length allowed by a model. We then computed the mean of each chunk embedding across the sequence dimension, and then averaged the mean embedding of each chunk to obtain the final embedding.

**Fig. A3:**
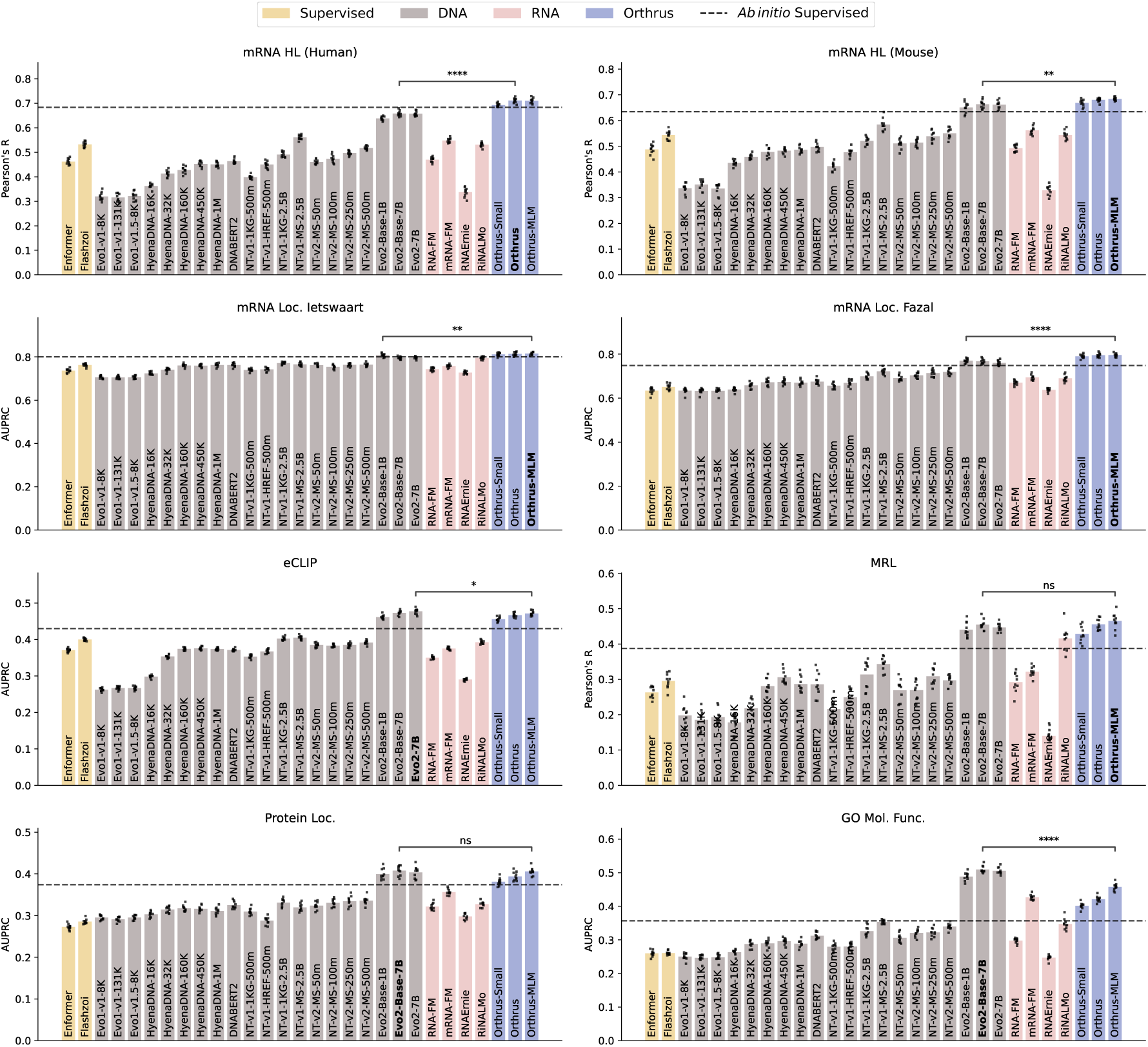
Benchmarking linear probing performance on mRNA property prediction tasks for self-supervised genomic foundation models with extended model variants. Error bars show 95% confidence intervals, constructed using 10 runs with randomized data splits. The grey dashed line indicates the performance of the maximum between Saluki and Dilated CNN models to better approximate real world performance

**Fig. A4:**
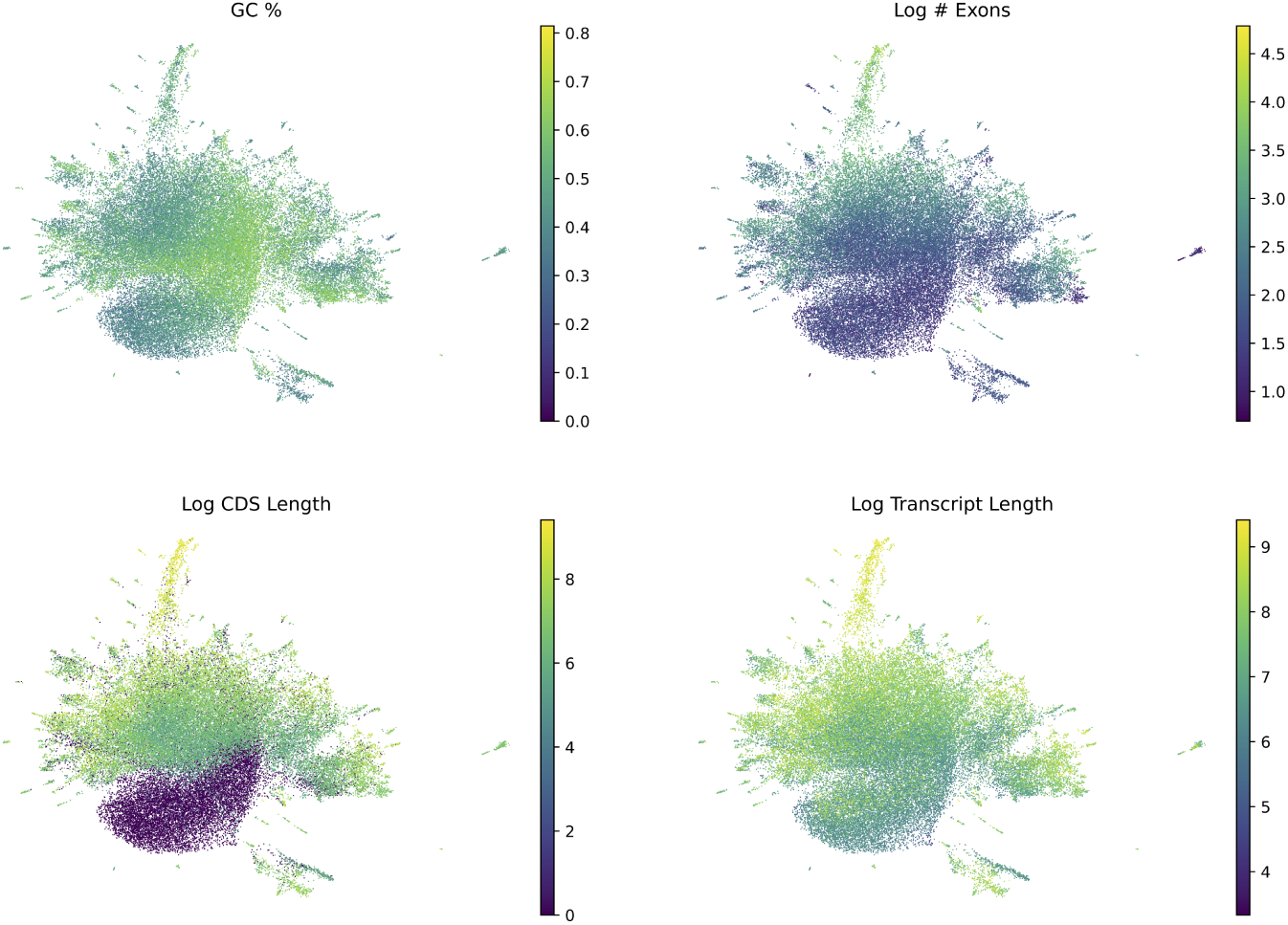
UMAP visualization of the Orthrus embeddings. 50,000 transcripts are randomly sampled from gencode comprehensive and colored according to transcript length, number of exons, CDS length, and GC%.

After obtaining embedding vectors, we used the scikit-learn implementation of linear models to perform the linear probes of the embeddings. For the downstream regression tasks, we used a linear

### A.5 Homology Splitting

To perform homology splitting we first acquire paralog information from Ensembl for a species of interest [76]. Ensembl provides pairs paralog information in the form of gene pairs related through duplication events. However, to perform homology splitting between genes we want to make sure that paralog transitivity is taken into account when dividing training samples between train, validation, and test splits. For example given three genes *g*_1_*, g*_2_*, g*_3_ if *g*_1_*, g*_2_ are annotated as paralogs and *g*_2_*, g*_3_ are annotated as well we want to ensure that *g*_1_*, g*_3_ are in the same split. Thus, we first transform the pairwise relationships into a graph structure in the process pruning low confidence paralog relationships. We enforce a similarity threshold of 35% which empirically demonstrated highly connected groups of paralogs. The algorithm for grouping is described in [27].

Following construction of the homology graph, during train-val-test split samples associated with genes which are connected in the homology graph are indexed into the same split. This avoids information leakage due to homology relationships within a given species.

### A.6 Fine-tuning Experimental Details

We fine-tune Orthrus Small by initializing the Mamba-backbone model with weights from pretraining and adding a two layer projection head with a hidden layer size of 256. We don’t apply any weight decay to weights that were initialized from pre-training while the projection head has an l2 weight decay term of 1e-3 or 1e-4 depending on the task. We fine-tune on downstream tasks using the Adam optimizer with a learning rate of 1e-3, 3e-4, or 1e-4 depending on the task. The models are trained with two NVIDIA L40s GPUs in a mixed precision setting. Go Molecular Function, RNA HL Human, and RNA HL Mouse were trained with weight decay / learning rate of 1e-3, RNA localization (Fazal and Ietswaart) and eCLIP were trained with 1e-4, and MRL Sugimoto was trained with 1e-4 weight decay and 3e-4 learning rate. We re-implemented the Saluki architecture in PyTorch to serve as a baseline. This implementation achieved a score of 0.75 on the joint mouse and human prediction task, a slight decrease from the original published score of 0.77.

Algorithm 1 Homology Group Assignment

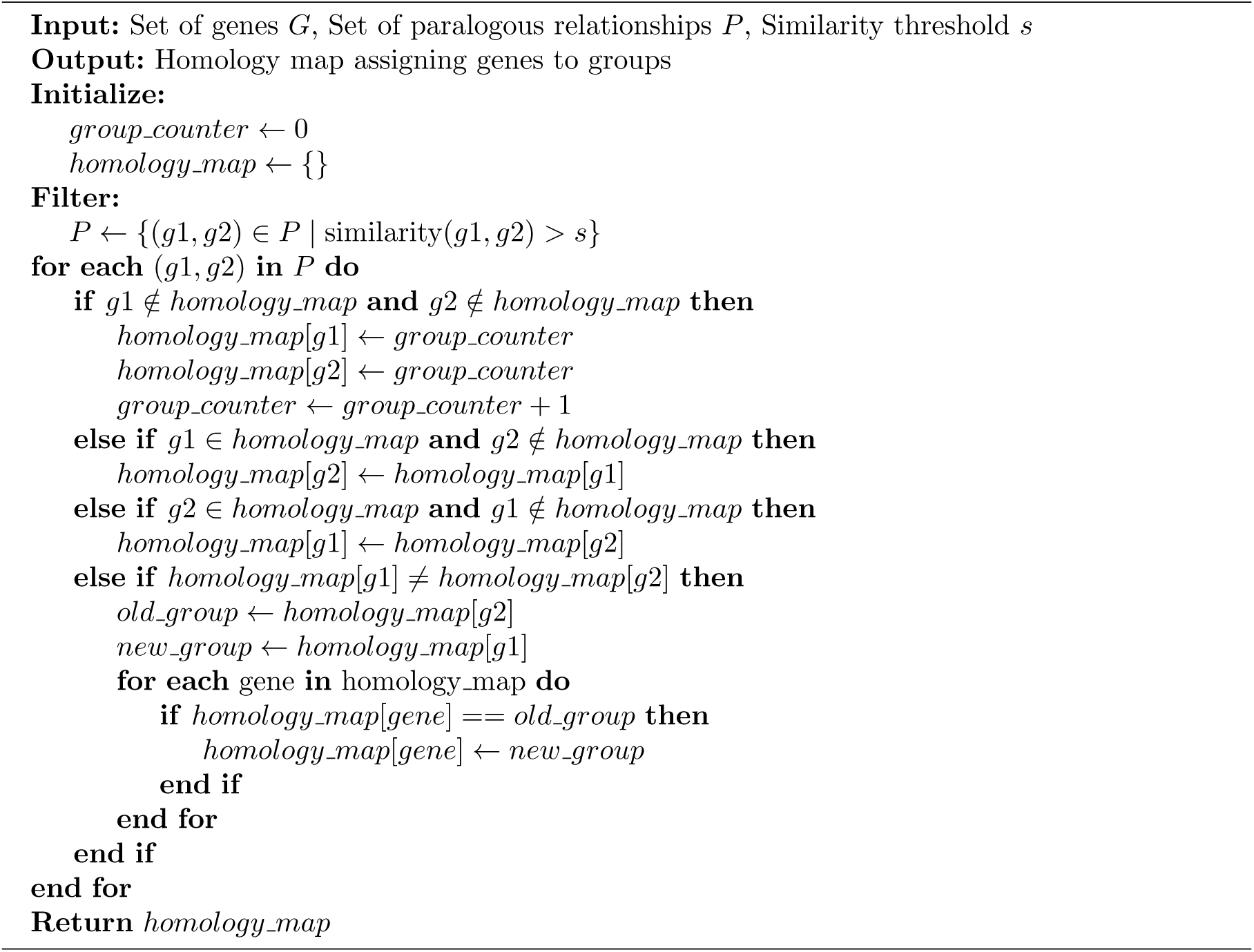

**Fig. A5:**
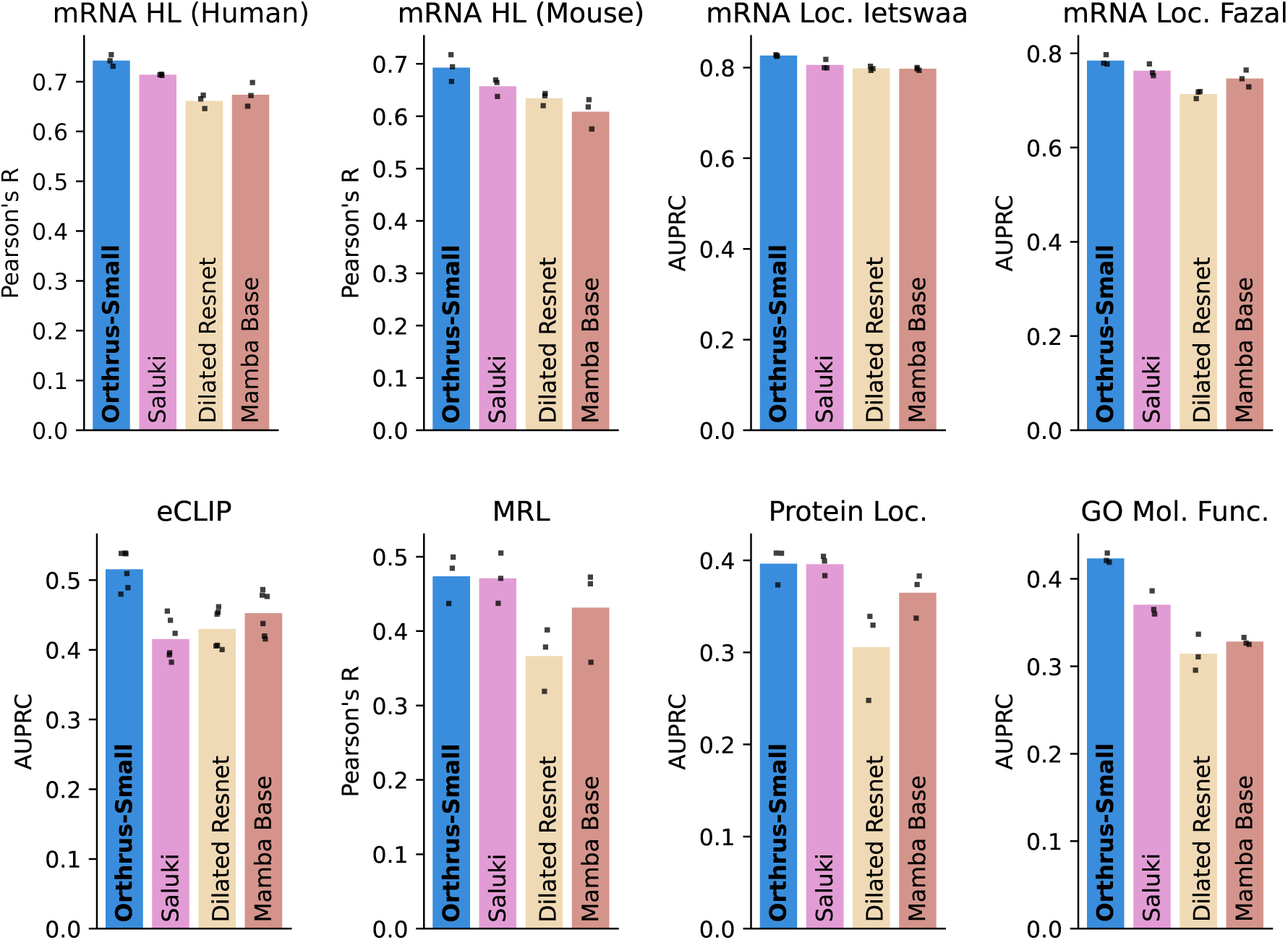
Fine tuning results plotted for 100% fraction of the data. Three seeds are trained per run and individual data points are visualized. Datasets are split with a homology aware strategy to avoid data leakage.

**Table A1:**
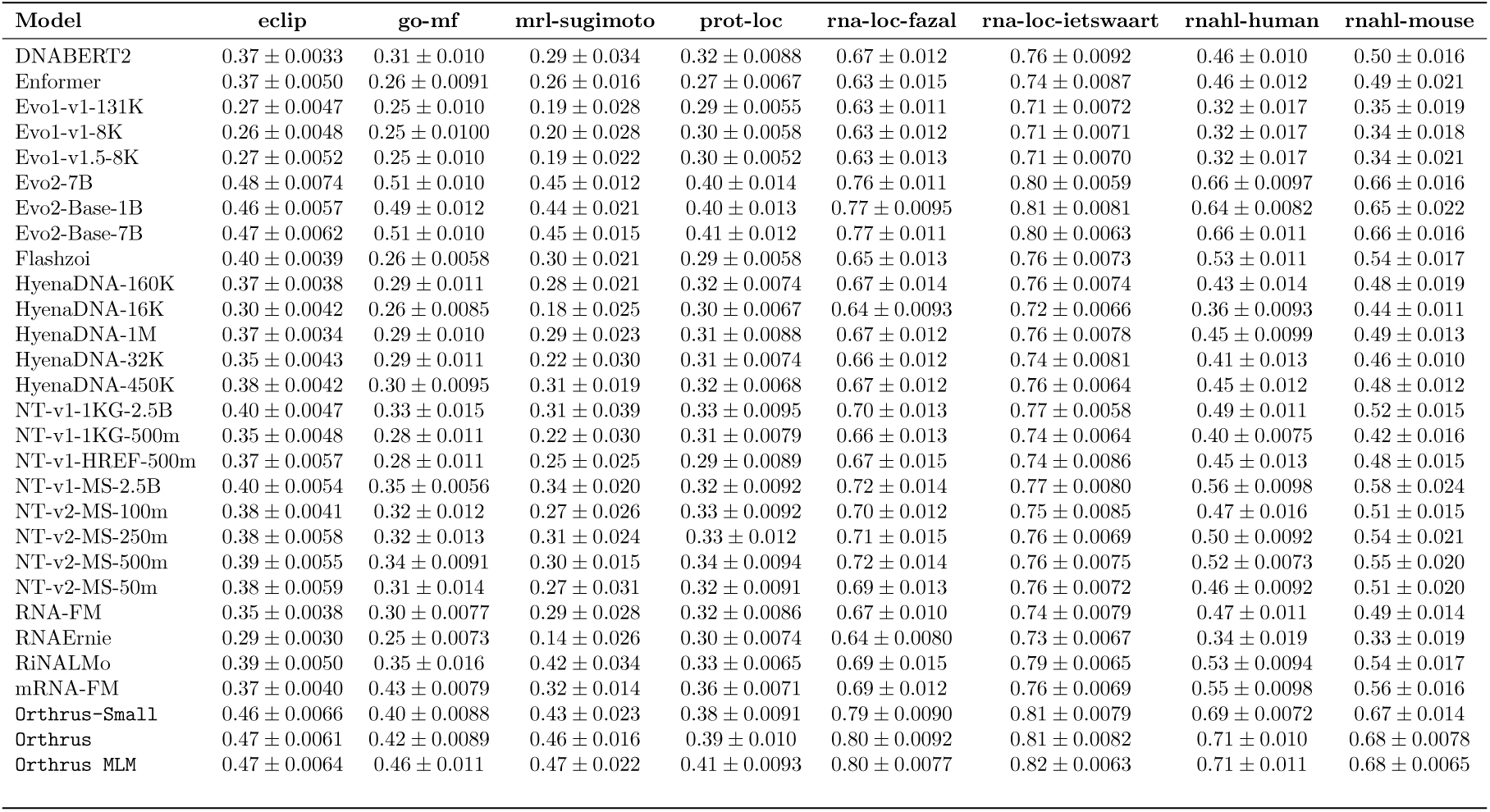
Linear probing results for self-supervised methods. The embeddings were computed for each method and then linear models were fit using the corresponding labels for each task. 95% Confidence intervals generated using 10 runs from random initialization.

**Table A2:** Pearson correlations (R) and auPRC of full model fine-tuning on mRNA half-life (HL), mean ribosome load (MRL), gene ontology molecular function (GO MF) classification, protein localization, RNA localization and eCLIP. Performance values were averaged over 3 random seeds. Training validation and test were split based on sequence homology to prevent data leakage.

| Dataset (Metric) | Orthrus Small | Dilated CNN | Mamba Small | Saluki |
| --- | --- | --- | --- | --- |
| eclip (auPRC) | $0.516 \pm 0.007$ | $0.430 \pm 0.001$ | $0.452 \pm 0.005$ | $0.415 \pm 0.005$ |
| go-mf (auPRC) | $0.423 \pm 0.006$ | $0.314 \pm 0.021$ | $0.328 \pm 0.004$ | $0.370 \pm 0.014$ |
| prot-loc (auPRC) | $0.396 \pm 0.023$ | $0.306 \pm 0.050$ | $0.365 \pm 0.024$ | $0.396 \pm 0.018$ |
| rna-loc-fazal (auPRC) | $0.784 \pm 0.011$ | $0.713 \pm 0.008$ | $0.747 \pm 0.018$ | $0.763 \pm 0.009$ |
| rna-loc-ietswaart (auPRC) | $0.826 \pm 0.002$ | $0.798 \pm 0.005$ | $0.797 \pm 0.003$ | $0.806 \pm 0.004$ |
| mrl-sugimoto (R) | $0.474 \pm 0.033$ | $0.366 \pm 0.043$ | $0.431 \pm 0.064$ | $0.471 \pm 0.040$ |
| rnahl-human (R) | $0.742 \pm 0.012$ | $0.661 \pm 0.014$ | $0.674 \pm 0.024$ | $0.713 \pm 0.019$ |
| rnahl-mouse (R) | $0.693 \pm 0.025$ | $0.634 \pm 0.012$ | $0.608 \pm 0.029$ | $0.657 \pm 0.030$ |

## Footnotes

1 In the coding region (2% of human DNA), an average individual carries 27 ± 13 unique SNPs [23].

## References

[1] Xiong, H. Y. et al. The human splicing code reveals new insights into the genetic determinants of disease. Science 347 (2015). URL 10.1126/science.1254806.

[2] Van Nostrand, E. L. et al. Robust transcriptome-wide discovery of RNA-binding protein binding sites with enhanced CLIP (eCLIP). Nat. Methods 13, 508–514 (2016).

[3] Brar, G. A. & Weissman, J. S. Ribosome profiling reveals the what, when, where and how of protein synthesis. Nature reviews Molecular cell biology 16, 651–664 (2015).

[4] Herzog, V. A. et al. Thiol-linked alkylation of rna to assess expression dynamics. Nature methods 14, 1198–1204 (2017).

[5] Jaganathan, K. et al. Predicting Splicing from Primary Sequence with Deep Learning. Cell 176, 535–548 (2019).

[6] Linder, J., Koplik, S. E., Kundaje, A. & Seelig, G. Deciphering the impact of genetic variation on human polyadenylation using APARENT2. Genome Biol 23, 232 (2022).

[7] Agarwal, V. & Kelley, D. R. The genetic and biochemical determinants of mRNA degradation rates in mammals. Genome Biol 23, 245 (2022).

[8] Merico, D. et al. G p.Met645Arg causes Wilson disease by promoting exon 6 skipping. NPJ Genom Med 5, 16 (2020).

[9] Richards, S. et al. Standards and guidelines for the interpretation of sequence variants: a joint consensus recommendation of the American College of Medical Genetics and Genomics and the Association for Molecular Pathology. Genet Med 17, 405–424 (2015).

10. Celaj, A., et al. An rna foundation model enables discovery of disease mechanisms and candidate therapeutics. *bioRxiv* (2023).

[11] Lotfollahi, M. et al. Predicting cellular responses to complex perturbations in high-throughput screens. Molecular Systems Biology 19 (2023).

[12] Ji, Y., Zhou, Z., Liu, H. & Davuluri, R. V. DNABERT: pre-trained Bidirectional Encoder Representations from Transformers model for DNA-language in genome. Bioinformatics 37, 2112–2120 (2021).

13. Chen, J., et al. Interpretable RNA Foundation Model from Unannotated Data for Highly Accurate RNA Structure and Function Predictions. *arXiv e-prints* arXiv:2204.00300 (2022).

[14] Nguyen, E., et al. HyenaDNA: Long-Range Genomic Sequence Modeling at Single Nucleotide Resolution. *arXiv e-prints* (2023).

[15] Peníc, R. J., Vlsíc, T., Huber, R. G., Wan, Y. & S^̌^ikíc, M. Rinalmo: general-purpose rna language models can generalize well on structure prediction tasks. Nature Communications 16 (2025). URL 10.1038/s41467-025-60872-5.

[16] Li, S. et al. mrna-lm: full-length integrated slm for mrna analysis. Nucleic Acids Research 53 (2025). URL 10.1093/nar/gkaf044.

[17] Nguyen, E. et al. Sequence modeling and design from molecular to genome scale with evo. Science 386 (2024). URL 10.1126/science.ado9336.

[18] Yuan, Y., Chen, Q. & Pan, X. Dgrna: a long-context rna foundation model with bidirectional attention mamba2 (2024). URL 10.1101/2024.10.31.621427.

[19] Brixi, G. et al. Genome modeling and design across all domains of life with evo 2 (2025). URL 10.1101/2025.02.18.638918.

[20] Devlin, J., Chang, M.-W., Lee, K. & Toutanova, K. BERT: Pre-training of Deep Bidirectional Transformers for Language Understanding. arXiv e-prints arXiv:1810.04805 (2018).

21. Radford, A. Improving language understanding by generative pre-training (2018).

[22] Song, Y., Wang, T., Mondal, S. K. & Sahoo, J. P. A comprehensive survey of few-shot learning: Evolution, applications, challenges, and opportunities (2022). URL https://arxiv.org/abs/2205. 06743. 2205.06743.

[23] Gudmundsson, S. et al. Variant interpretation using population databases: Lessons from gnomAD. Human Mutation 43, 1012–1030 (2021).

[24] Taliun, D. et al. Sequencing of 53,831 diverse genomes from the NHLBI TOPMed Program. Nature 590, 290–299 (2021).

[25] Chen, S. et al. A genomic mutational constraint map using variation in 76, 156 human genomes. Nature 625, 92–100 (2023). URL 10.1038/s41586-023-06045-0.

[26] Lindblad-Toh, K. et al. A high-resolution map of human evolutionary constraint using 29 mammals. Nature 478, 476–482 (2011).

[27] Sullivan, P. F. et al. Leveraging base-pair mammalian constraint to understand genetic variation and human disease. Science 380 (2023). URL 10.1126/science.abn2937.

[28] Christmas, M. J. et al. Evolutionary constraint and innovation across hundreds of placental mammals. Science 380 (2023). URL 10.1126/science.abn3943.

[29] Dalla-Torre, H., et al. The nucleotide transformer: Building and evaluating robust foundation models for human genomics. *bioRxiv* (2023).

[30] Lu, A. X., Lu, A. X. & Moses, A. Evolution Is All You Need: Phylogenetic Augmentation for Contrastive Learning. arXiv e-prints (2020).

31. Chen, T., Kornblith, S., Norouzi, M. & Hinton, G. A Simple Framework for Contrastive Learning of Visual Representations. *arXiv e-prints* arXiv:2002.05709 (2020).

[32] Kirilenko, B. M. et al. Integrating gene annotation with orthology inference at scale. Science 380 (2023). URL 10.1126/science.abn3107.

[33] Gu, A. & Dao, T. Mamba: Linear-time sequence modeling with selective state spaces (2024). 2312.00752.

[34] Fair, B. et al. Global impact of unproductive splicing on human gene expression. Nature Genetics 56, 1851–1861 (2024). URL 10.1038/s41588-024-01872-x.

[35] Schertzer, M. D. et al. Cas13d-mediated isoform-specific rna knockdown with a unified computational and experimental toolbox. bioRxiv 2023.09.12.557474 (2023). URL https://www.biorxiv.org/content/10.1101/2023.09.12.557474v1.

[36] Koonin, E. V. Orthologs, paralogs, and evolutionary genomics. Annual Review of Genetics **39**, 309–338 (2005). URL 10.1146/annurev.genet.39.073003.114725.

37. McInnes, L., Healy, J. & Melville, J. Umap: Uniform manifold approximation and projection for dimension reduction (2020). URL https://arxiv.org/abs/1802.03426. 1802.03426.

[38] Frankish, A. et al. GENCODE 2021. Nucleic Acids Res 49, D916–D923 (2021).

[39] O’Leary, N. A. et al. Reference sequence (RefSeq) database at NCBI: current status, taxonomic expansion, and functional annotation. Nucleic Acids Res 44, D733–745 (2016).

[40] Koonin, E. V. & Wolf, Y. I. Constraints and plasticity in genome and molecular-phenome evolution. Nature Reviews Genetics 11, 487–498 (2010). URL 10.1038/ nrg2810.

[41] Sayers, E. W. et al. Database resources of the National Center for Biotechnology Information in 2023. Nucleic Acids Res 51, D29–D38 (2023).

42. Yeh, C.-H., et al. Decoupled Contrastive Learning. *arXiv e-prints* arXiv:2110.06848 (2021).

[43] Kelley, D. R. et al. Sequential regulatory activity prediction across chromosomes with convolutional neural networks. Genome Res 28, 739–750 (2018).

[44] Karollus, A. et al. Species-aware dna language models capture regulatory elements and their evolution. Genome Biology 25 (2024). URL 10.1186/s13059-024-03221-x.

[45] Abramson, J. et al. Accurate structure prediction of biomolecular interactions with alphafold 3. Nature 630, 493–500 (2024). URL 10.1038/s41586-024-07487-w.

46. Schrodinger, L. & DeLano, W. Pymol. URL http://www.pymol.org/pymol.

[47] Warren, C. F. A., Wong-Brown, M. W. & Bowden, N. A. BCL-2 family isoforms in apoptosis and cancer. Cell Death Dis. 10, 177 (2019).

[48] Loo, L. S. W. et al. Bcl-xl/bcl2l1 is a critical anti-apoptotic protein that promotes the survival of differentiating pancreatic cells from human pluripotent stem cells. Cell Death Disease 11 (2020). URL 10.1038/s41419-020-2589-7.

[49] Wickenhagen, A. et al. A prenylated dsRNA sensor protects against severe COVID-19. Science 374, eabj3624 (2021).

[50] Lee, N. K., Tang, Z., Toneyan, S. & Koo, P. K. EvoAug: improving generalization and interpretability of genomic deep neural networks with evolution-inspired data augmentations. Genome Biol 24, 105 (2023).

[51] Lu, A. X., Zhang, H., Ghassemi, M. & Moses, A. Self-supervised contrastive learning of protein representations by mutual information maximization. bioRxiv (2020).

[52] Pertea, M. et al. CHESS: a new human gene catalog curated from thousands of large-scale RNA sequencing experiments reveals extensive transcriptional noise. Genome Biol 19, 208 (2018).

[53] von Kügelgen, J., et al. Self-supervised learning with data augmentations provably isolates content from style (2022). 2106.04619.

[54] Xiao, M.-S. et al. Genome-scale exon perturbation screens uncover exons critical for cell fitness. Molecular Cell 84, 2553–2572.e19 (2024). URL 10.1016/j.molcel.2024.05.024.

[55] Spies, N., Burge, C. B. & Bartel, D. P. 3’ UTR-isoform choice has limited influence on the stability and translational efficiency of most mRNAs in mouse fibroblasts. Genome Res 23, 2078–2090 (2013).

[56] Irimia, M. et al. A highly conserved program of neuronal microexons is misregulated in autistic brains. Cell 159, 1511–1523 (2014).

[57] Garćıa-Perez, R., et al. The landscape of expression and alternative splicing variation across human traits. Cell Genomics 0 (2022).

[58] van den Oord, A., Li, Y. & Vinyals, O. Representation Learning with Contrastive Predictive Coding. arXiv e-prints (2018).

[59] Vaswani, A. Attention is all you need. Advances in Neural Information Processing Systems (2017).

60. Gu, A., Goel, K. & Ŕe, C. Efficiently modeling long sequences with structured state spaces (2022). URL https://arxiv.org/abs/2111.00396. 2111.00396.

61. Georgakopoulos-Soares, I., et al. Transcription factor binding site orientation and order are major drivers of gene regulatory activity. *Nature Communications* 14 (2023). URL http://dx.doi.org/10.1038/s41467-023-37960-5.

[62] Sohn, K. Lee, D., Sugiyama, M., Luxburg, U., Guyon, I. & Garnett, R. (eds) *Improved deep metric learning with multi-class n-pair loss objective*. (eds Lee, D., Sugiyama, M., Luxburg, U., Guyon, I. & Garnett, R.) *Advances in Neural Information Processing Systems*, Vol. 29 (Curran Associates, Inc., 2016).

[63] Sugimoto, Y. & Ratcliffe, P. J. Isoform-resolved mRNA profiling of ribosome load defines interplay of HIF and mTOR dysregulation in kidney cancer. Nature Structural Molecular Biology 29, 871–880 (2022).

[64] Thul, P. J. et al. A subcellular map of the human proteome. Science 356 (2017).

[65] Rodriguez, J. M. et al. APPRIS: selecting functionally important isoforms. Nucleic Acids Res. 50, D54–D59 (2022).

[66] Consortium, T. G. O. et al. The Gene Ontology knowledgebase in 2023. Genetics 224, iyad031 (2023).

[67] Ashburner, M. et al. Gene ontology: tool for the unification of biology. Nature Genetics 25, 25–29 (2000).

[68] Zhou, N. et al. The CAFA challenge reports improved protein function prediction and new functional annotations for hundreds of genes through experimental screens. Genome Biol 20, 244 (2019).

[69] Rodriguez, J. M. et al. Appris: annotation of principal and alternative splice isoforms. Nucleic acids research 41, D110–D117 (2013).

[70] Shi, R., et al. mrnabench: A curated benchmark for mature mrna property and function prediction (2024). Unpublished manuscript.

[71] Consortium, E. P. et al. An integrated encyclopedia of dna elements in the human genome. Nature 489, 57 (2012).

[72] Van Nostrand, E. L. et al. Robust transcriptome-wide discovery of rna-binding protein binding sites with enhanced clip (eclip). Nature methods 13, 508–514 (2016).

[73] Ietswaart, R. et al. Genome-wide quantification of rna flow across subcellular compartments reveals determinants of the mammalian transcript life cycle. Molecular Cell 84, 2765–2784 (2024).

[74] Fazal, F. M. et al. Atlas of subcellular rna localization revealed by apex-seq. Cell 178, 473–490 (2019).

[75] Jumper, J. et al. Highly accurate protein structure prediction with AlphaFold. Nature 596, 583–589 (2021).

[76] Martin, F. J. et al. Ensembl 2023. Nucleic Acids Research 51, D933–D941 (2022). URL 10.1093/nar/gkac958.

[77] Zar, J. H. *Biostatistical Analysis* 5 edn (Pearson, Upper Saddle River, NJ, 2009).

[78] Zhang, Z. et al. Protein language models learn evolutionary statistics of interacting sequence motifs. Proceedings of the National Academy of Sciences 121 (2024). URL 10.1073/pnas.2406285121.

[79] Tomaz da Silva, P., et al. Nucleotide dependency analysis of dna language models reveals genomic functional elements (2024). URL 10.1101/2024.07.27.605418.

[80] Avsec, Z. et al. Effective gene expression prediction from sequence by integrating longrange interactions. Nature Methods 18, 1196–1203 (2021). URL 10.1038/ s41592-021-01252-x.

[81] Linder, J., Srivastava, D., Yuan, H., Agarwal, V. & Kelley, D. R. Predicting rna-seq coverage from dna sequence as a unifying model of gene regulation. Nature Genetics 57, 949–961 (2025). URL 10.1038/s41588-024-02053-6.

[82] Hingerl, J. C., Karollus, A. & Gagneur, J. Flashzoi: An enhanced borzoi model for accelerated genomic analysis (2024). URL 10.1101/2024.12.18.629121.

